# *Leishmania* infection induces a limited differential gene expression in the sand fly midgut

**DOI:** 10.1101/845867

**Authors:** Iliano V. Coutinho-Abreu, Tiago D. Serafim, Claudio Meneses, Shaden Kamhawi, Fabiano Oliveira, Jesus G. Valenzuela

## Abstract

**Background:** Phlebotomine sand flies are the vectors of *Leishmania* worldwide. To develop in the sand fly midgut, *Leishmania* multiplies and undergoes multiple stage differentiations leading to the infective form, the metacyclic promastigotes. To gain a better understanding of the influence of *Leishmania* infection in midgut gene expression, we performed RNA-Seq comparing uninfected *Lutzomyia longipalpis* midguts and *Leishmania infantum*-infected *Lutzomyia longipalpis* midguts at seven time points which cover the various developmental *Leishmania* stages including early time points when blood digestion is taking place and late time points when the parasites are undergoing metacyclogenesis.

**Results:** Out of over 13,841 transcripts assembled *de novo*, only 113 sand fly transcripts, about 1%, were differentially expressed. Further, we observed a low overlap of differentially expressed sand fly transcripts across different time points suggesting a specific influence of each *Leishmania* stage on midgut gene expression. Two main patterns of sand fly gene expression modulation were noticed. At early time points (days 1-4), more transcripts were down-regulated by *Leishmania* infection at large fold changes (> −32 fold). Among the down-regulated genes, the transcription factor Forkhead/HNF-3 and hormone degradation enzymes were differentially regulated on day 4 and appear to be the upstream regulators of nutrient transport, digestive enzymes, and peritrophic matrix proteins. Conversely, at later time points (days 6 onwards), most of the differentially expressed transcripts were up-regulated by small fold changes (< 32 fold), and the molecular function of such genes are associated with the metabolism of lipids and detoxification of xenobiotics (P450).

**Conclusion:** Overall, it appears that *Leishmania* modulates sand fly gene expression early on in order to overcome the barriers imposed by the midgut, yet it behaves like a commensal at later time points, when modest midgut gene expression changes correlate with a massive amount of parasites in the anterior midgut.

## Background

*Leishmania* is a digenetic parasite developing in the mammalian host as well as in the insect vector. These parasites are mostly transmitted by phlebotomine sand flies (Diptera: Psychodidae) of the genera *Phlebotomus* and *Lutzomyia* in the Old and New World, respectively [1].

*Leishmania* fully develops in the lumen of the sand fly midgut [2–4]. Once a sand fly takes up an infected blood meal, *Leishmania* is carried along within macrophages in the round-shaped amastigote form (mammalian stage). Between 18h and 24h post blood meal, these parasites are released from the macrophages and start to differentiate into procyclic promastigotes within blood enveloped by the peritrophic matrix [5]. During this process, the parasites elongate their cell bodies and expose their flagella, becoming fully differentiated into procyclics by day 2 (48h). Between days 2 and 4, *Leishmania* multiplies and undergoes another differentiation step, acquiring an elongated (banana-like shape) form termed nectomonads [2–4]. Upon the breakdown of the peritrophic matrix, the nectomonads escape to the ectoperitrophic space and eventually dock on the midgut microvilli [6, 7]. As the remains of the digested blood are evacuated, the parasites detach from the epithelium and further differentiate into the leptomonad stage, which exhibit a smaller cell body and a longer flagellum than nectomonads [2–4]. From day 6 onwards, the leptomonads undergo a differentiation process, termed metacyclogenesis, giving rise to the infective forms, the metacyclic promastigotes [8]. During metacyclogenesis, the parasites replace their glycocalyx, exhibiting different sugar side chains on their major surface glycans, reduce the size of their cell bodies, and elongate their flagella [2–4]. All these transformations give rise to highly motile parasites [2–4].

Even when developing in their natural sand fly vectors, *Leishmania* faces barriers imposed by the midgut; overtaking such barriers is critical for the development of mature *Leishmania* infections. During the transitional stages between amastigotes and procyclic promastigotes, the parasites are susceptible to the harmful action of digestive enzymes [9]. The immune system may also counteract infection with the parasites, by activation of the Imd pathway [10, 11]. Escaping from the peritrophic matrix is also a crucial step for *Leishmania* survival [12, 13]. Another critical barrier is the attachment to the midgut epithelium [14]. For this step, specific carbohydrate side chains are required for binding to a midgut epithelium receptor [7, 15, 16]. From there on, undefined parameters trigger the metacyclogenesis process in parasites leading to the development of a mature infection.

The midgut transcriptomes of three sand fly species have been described, focusing mostly on differences in gene expression triggered by blood intake and parasite infection as compared to sugar fed midguts [18–20]. Nonetheless, such studies took place before the advent of deep sequencing, being limited to the investigation of about 1,000 transcripts due to the low dynamic range of cDNA libraries. Despite such a limited pool of genes, these studies unveiled multiple genes differentially regulated by blood and/or *Leishmania* infection. For the later, genes encoding digestive enzymes and components of the peritrophic matrix, the main midgut barriers to *Leishmania* development, were differentially regulated [18–20].

In order to investigate the effects of *Leishmania* infection on sand fly midgut gene expression, we carried out an RNA-Seq analysis of *Leishmania infantum*-infected *Lutzomyia longipalpis* midguts at 7 timepoints, each corresponding to when the insect midguts are enriched with a particular *Leishmania* stage. These encompassed early time points when blood digestion is taking place as well as late time points when the parasites are undergoing metacyclogenesis. This approach expands our breadth of knowledge by assessing the effects of *Leishmania* infection on over 13,000 sand fly midgut transcripts, focusing on genes encoding secreted proteins and also on genes participating in biological processes.

## Results

### Sand fly infection and *Leishmania* differentiation

In order to assess how gene expression in sand fly midguts is affected by *Leishmania* growth and differentiation, *Le. infantum* infected-*Lutzomyia longipalpis* midguts (1d through 14d Pi) were dissected for RNA-Seq library construction in triplicate and compared to midguts fed on uninfected blood at the same time points (1d through 14d PBM). All the libraries gave rise to high quality data and robust expression levels, except one replicate of the 2d PBM and another of the 12d PBM time points, which were excluded from further analyses. For infected midguts, *Le. infantum* growth in the *Lu. longipalpis* sand fly midgut followed a typical and expected pattern whereby low levels of parasites were detected early at 4d (median = 3,000 parasites) and 6d (median = 6,000 parasites). From 6d to 14d, the parasite load increased 21-fold, reaching about 126,000 parasites at 14d. During the late time points, parasites underwent differentiation through. metacyclogenesis, increasing the proportion of metacyclic stage parasites from 0% on 6d to 92% on 14d [21].

### Expanding the *Lu. longipalpis* midgut repertoire of putative proteins

A *Lu. longipalpis* midgut-specific *de novo* assembly was made from libraries prepared from RNA extracted from uninfected midguts at each of the seven study timepoints (total of 53,683,499 high quality reads). High quality reads were assembled in 57,016 contigs that were further down-selected to 13,841 putative contigs based on the presence of an ORF and similarities to proteins deposited at Refseq invertebrate, NCBI Genbank or SwissProt. Putative proteins where a signal peptide was predicted were also considered. Selected contigs varied in size with the shortest at 150 bp, the longest at 27,627 bp and the mean size at 1,498bp. Overall, 72% could be categorized to a functional class after BLAST analysis (e<10E-6) to nine distinct databases (Additional file 1: Fig. S1 and Additional file 2: Table S1). Figure 1 shows an overview of the transcriptome repertoire displaying the overall percentage of contigs (% of contigs) or abundance as transcripts per million (%TPM) for all time points and conditions combined, highlighting the distribution of the mapped reads to the functional classification. Unknown contigs accounted for 28% of the contigs, but only for 6.56 % of transcriptome abundance. The most represented functional categories were secreted proteins with 25.9 % of TPM, protein synthesis (15.2 % of TMP), metabolism (14.3 % of TMP) and protein modification (11.8 % of TMP) (Fig. 1 and Additional file 3: Table S2 and Additional file 4: Fig. S2). This dataset was used to map the individual samples and determine the sand fly midgut differential expression caused by *Leishmania* infection (Additional file 3: Table S2.)

**Figure 1.**
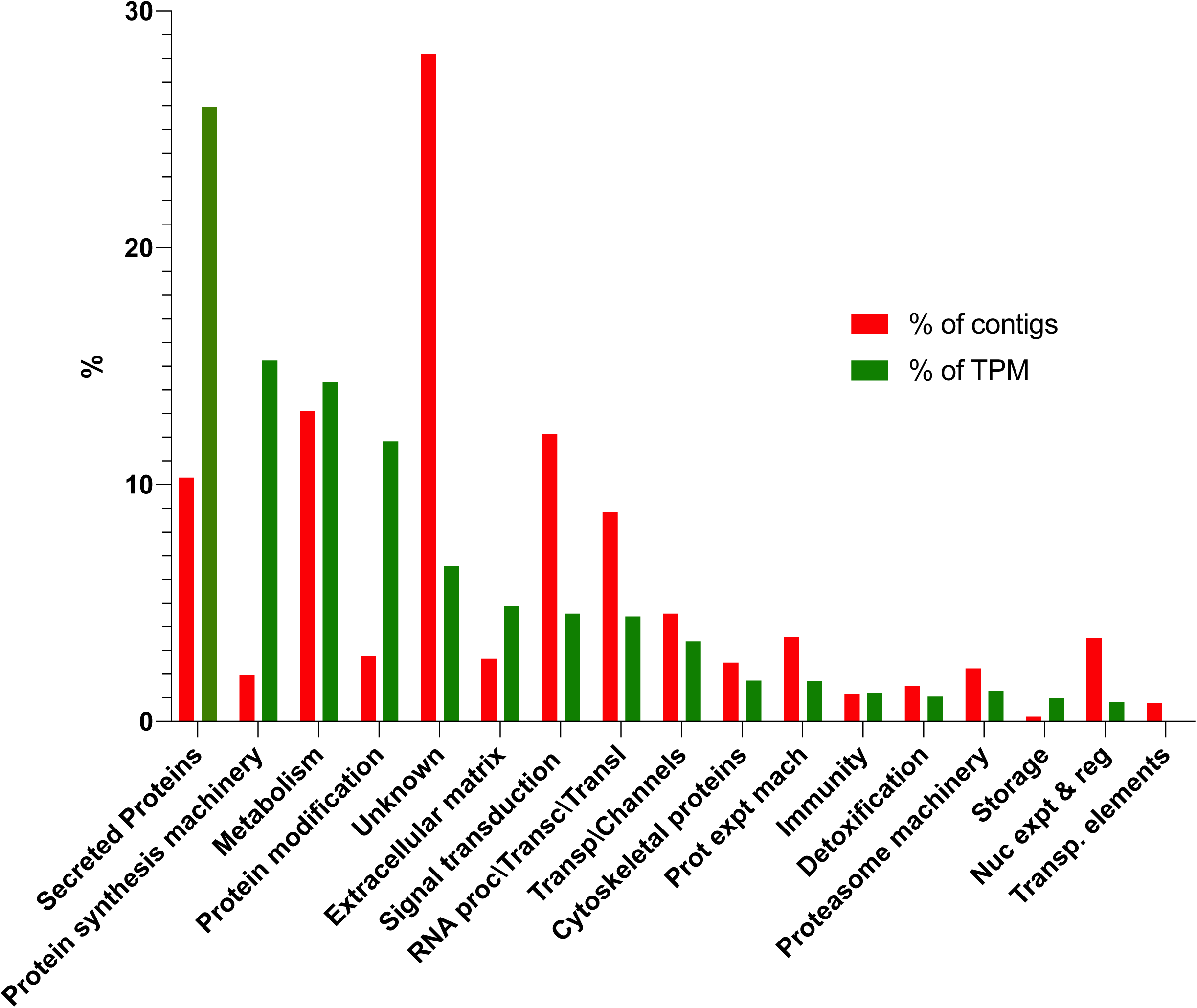
Overview of the transcriptome repertoire displaying the overall percentage of contigs (% of contigs) or abundance (%TPM) for all time points. The distribution of the mapped reads to the functional classification are highlighted.

### Sand fly midgut gene expression

The overall expression profiles of the infected and uninfected midguts obtained at seven time points each representing infected midguts enriched with a different *Leishmania* stage is summarized by PCA analyses of the average expression for each time point (Fig 2A and Additional file 5: Table S3) as well as amongst replicates (Additional file 6: Fig. S3 and Additional file 5: Table S3). The PC1 axis showed a clear separation between the midguts in which blood digestion is ongoing (Fig. 2A left side, 1d PBM/Pi and 2d PBM/Pi) from the time points at which the blood was mostly digested (Fig. 2A right side, 4d PBM/Pi) and the remaining time points where the midguts were clear of blood (Fig. 2A right side, 6d to 14 PBM/Pi). The PC1 accounted for 77.2% of the variance (Additional file 5: Table S3). On the other hand, the PC3 (rather than PC2; Additional file 6: Fig. S3B) sorted for the most part the infected from the uninfected samples (Fig. 2A) and only accounted for 4.1% of the variance (Additional file 5: Table S3).

**Figure 2.**
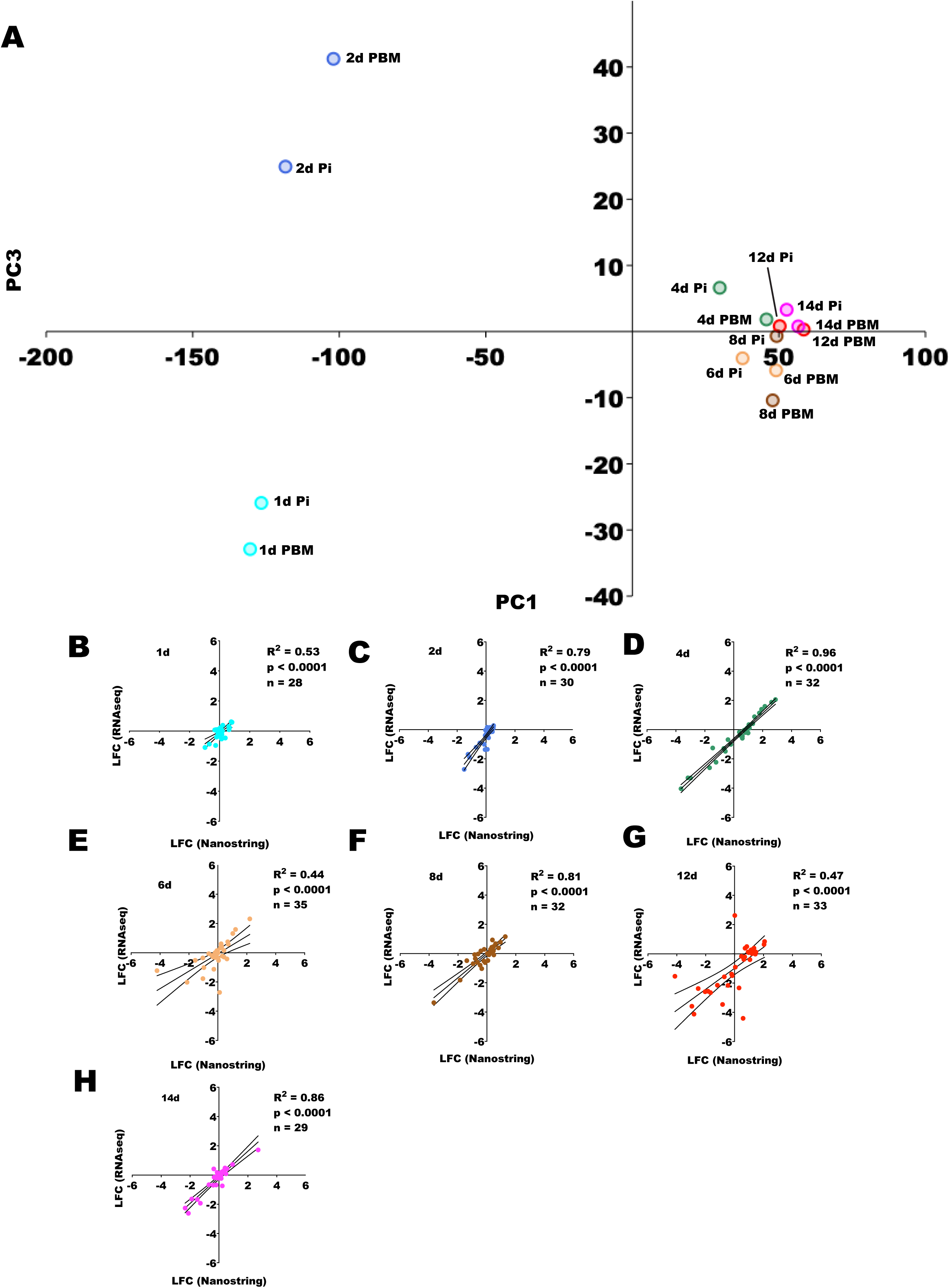
Midgut sequencing overall analysis. **A**. Principal component analysis (PCA) describing the position of each midgut time point on the expression space. Expression space was generated based on the log_2_ of TPMs using the 10,000 most highly expressed transcripts across libraries. The Eigenvalues and % variance for PC1 and PC3 % were 6221.99 and 77.19% and 330.34 and 4.1%, respectively. **B-H.** Gene expression validation by nCounter (Nanostring). Linear regression analyses comparing the expression profiles of randomly chosen transcripts obtained with RNA-Seq and nCounter (Nanostring) techniques for the seven time points. All comparisons were statistically significant (p < 0.0001). R^2^: regression coefficient. n: number of transcripts. The color codes labeling each time point were as follow: B. Aqua (1d); C. Royal Blue (2d); D. Sea Green (4d); E. Sandy Brown (6d); F. Saddle Brown (8d); G. Red (12d); and H. Fuchsia (14d).

The expression profiles of midguts were validated by assessing the expression levels of selected midgut genes (n = 28-35; Additional file 7: Table S4) using the nCounter technology (NanoString). The mean log_2_ fold change (LFC) of infected over uninfected samples was compared at each time point with LFC data obtained with the RNA-Seq technique for the same genes. Representative genes participate in chitin metabolism/ peritrophic matrix scaffolding (peritrophins and chitinases), immunity (defensin, catalase, and spatzle), digestion (amylase and chymotrypsin) among others are depicted in Fig. 2B-H. The regression analyses between the expression levels obtained with nCounter and RNA-Seq were statistically significant (p < 0.0001) for all seven time points (Fig. 2B-H), and the regression coefficients were greater than 0.5 for all time points, except 6d (R^2^ = 0.40) and 12d (R^2^ = 0.47) as shown in Fig. 2B-H.

### Modulation of sand fly midgut gene expression by *Leishmania* infection

Differences in gene expression between *Leishmania*-infected over uninfected midguts at the seven time points were assessed. Overall, such differences accounted for only 113 differentially expressed transcripts (1 < LFC > 1; q-value < 0.05; Additional file 8: Table S5). The number of DE genes gradually increased from 2 genes on 1d to 53 genes on 4d (Fig. 3A). On 6d, the number of DE genes decreased to 20 genes and went further down to 15 genes on 8d (Fig. 3A). Four days later, there was a strong increase in the number of DE genes (12d = 32 genes), which was reduced to 13 genes two days later at 14d (Fig. 3A).

**Figure 3.**
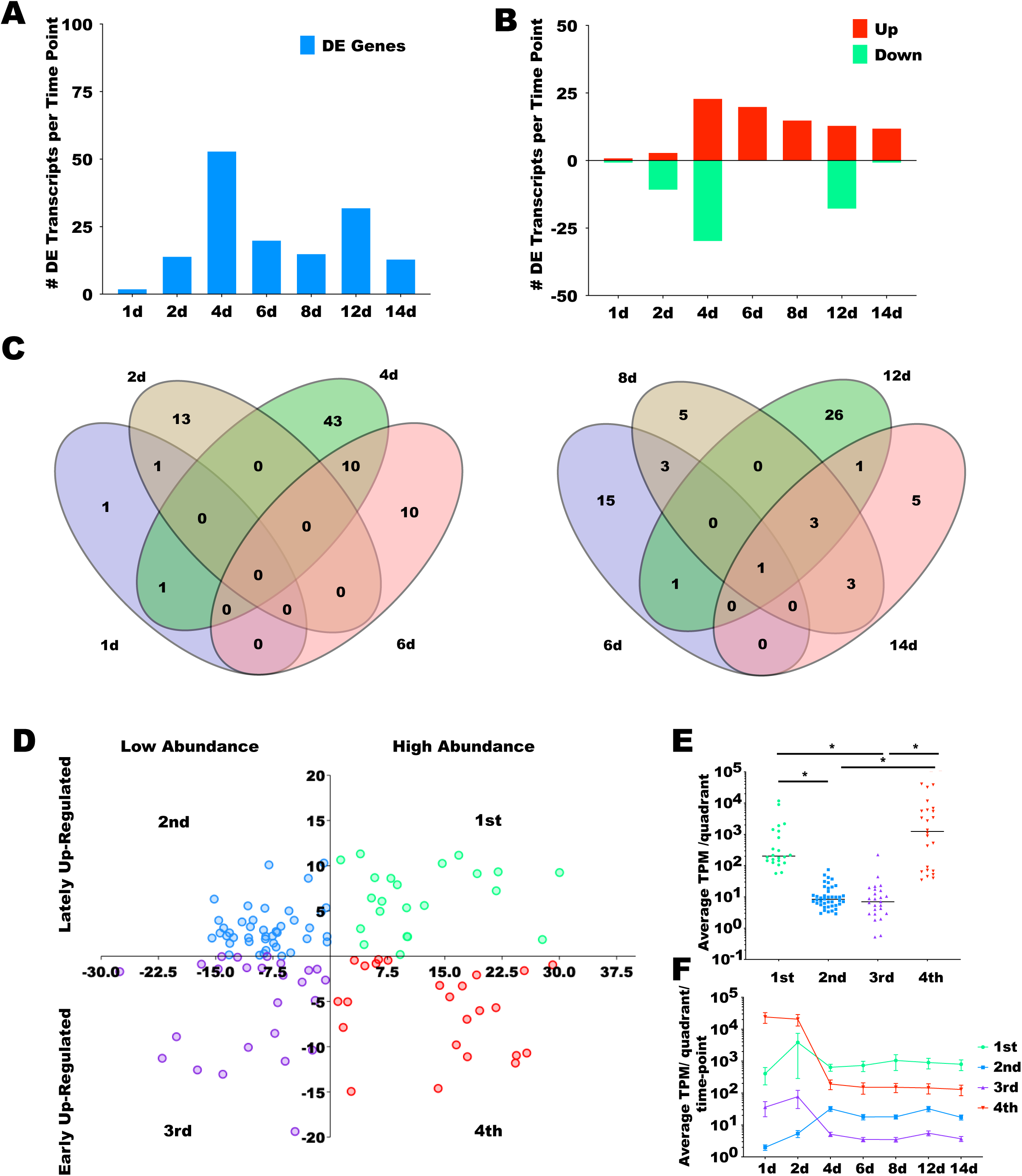
Analysis of differentially expressed (DE) midgut transcripts across time points. **A**. Total number of differentially expressed transcripts across time points. **B**. Number of DE transcripts up- and down-regulated in *Leishmania* infected over uninfected midguts at each time point. **C**. Left: Venn diagrams depicting the number of DE transcripts unique and shared amongst the time points 1d through 6d. Right: Venn diagrams depicting the number (and percentages) of DE transcripts unique and shared amongst time points 6d through 14d. **D**. PC analysis of all the DE transcripts in all time points based on the log_2_ fold change (LFC) of the *Leishmania*-infected over the uninfected TPM values for each transcript. Each quadrant in the expression space was labelled from 1^st^ to 4^th^ and the transcripts mapped to the respective quadrants were color coded in Spring Green (1^st^), Dodge Blue (2^nd^), Blue Violet (3^rd^), and Red (4^th^). The Eigenvalues and % variance for PC1 and PC2 % were 163.59 and 76.28% and 40.6 and 18.94%, respectively. **E**. Expression analysis per quadrant. The average TPM across time points for every DE transcript mapped onto each quadrant was plotted. Horizontal bars indicate median values and differences were statistically significant (* Mann Whitney U test, p < 0.0001). Color coding as in D. **F**. Expression analysis per quadrant per time point in blood fed libraries (PBM). The average TPM for each time point for every DE transcript mapped in each quadrant was plotted. Mean TPM as shapes and SEM bars are depicted. Based on the differences observed in E and F, the quadrants in D were labeled to describe the DE transcripts expressed in high and low abundance (as defined by PC1) and expressed early and late (as defined by PC2). DE was considered significant for transcripts displaying LFC either lower than −1 or higher than 1 and FDR q-value lower than 0.05.

Amongst the midgut genes differentially expressed upon *Leishmania* infection, some appear to play a role in specific biological processes (Table 1; Additional File 8: Table S5). A gene encoding the transcription factor Forkhead/HNF-3 (lulogut44569) was down-regulated on 2d. Genes encoding proteins potentially involved with metabolism of steroid hormones, such as 17-beta-hydroxysteroid dehydrogenase 13-like (lulogut32574) and juvenile hormone esterase (lulogut40195) were down-regulated on 2d; a putative juvenile hormone binding protein (lulogutSigP-24104) was down-regulated on 4d; and an ecdysteroid kinase (lulogut41307) was down-regulated on 12d. Also, genes encoding a peritrophic matrix protein (lulogutSigP-40401), involved with the peritrophic matrix scaffolding, the antimicrobial peptide attacin (lulogutSigP-8812), and amino acid (lulogut16004) and trehalose (lulogutSigP-40100) transporters, were down-regulated on 4d. Amongst the up-regulated genes, multiple peptidases and proteases were up-regulated on 4d and 6d. Likewise, multiple insect allergen proteins (microvilli proteins) of unknown function were up-regulated on 4d and 6d upon *Leishmania* infection. From 8d onwards, multiple cytochrome p450 transcripts were up-regulated.

**Table 1.**
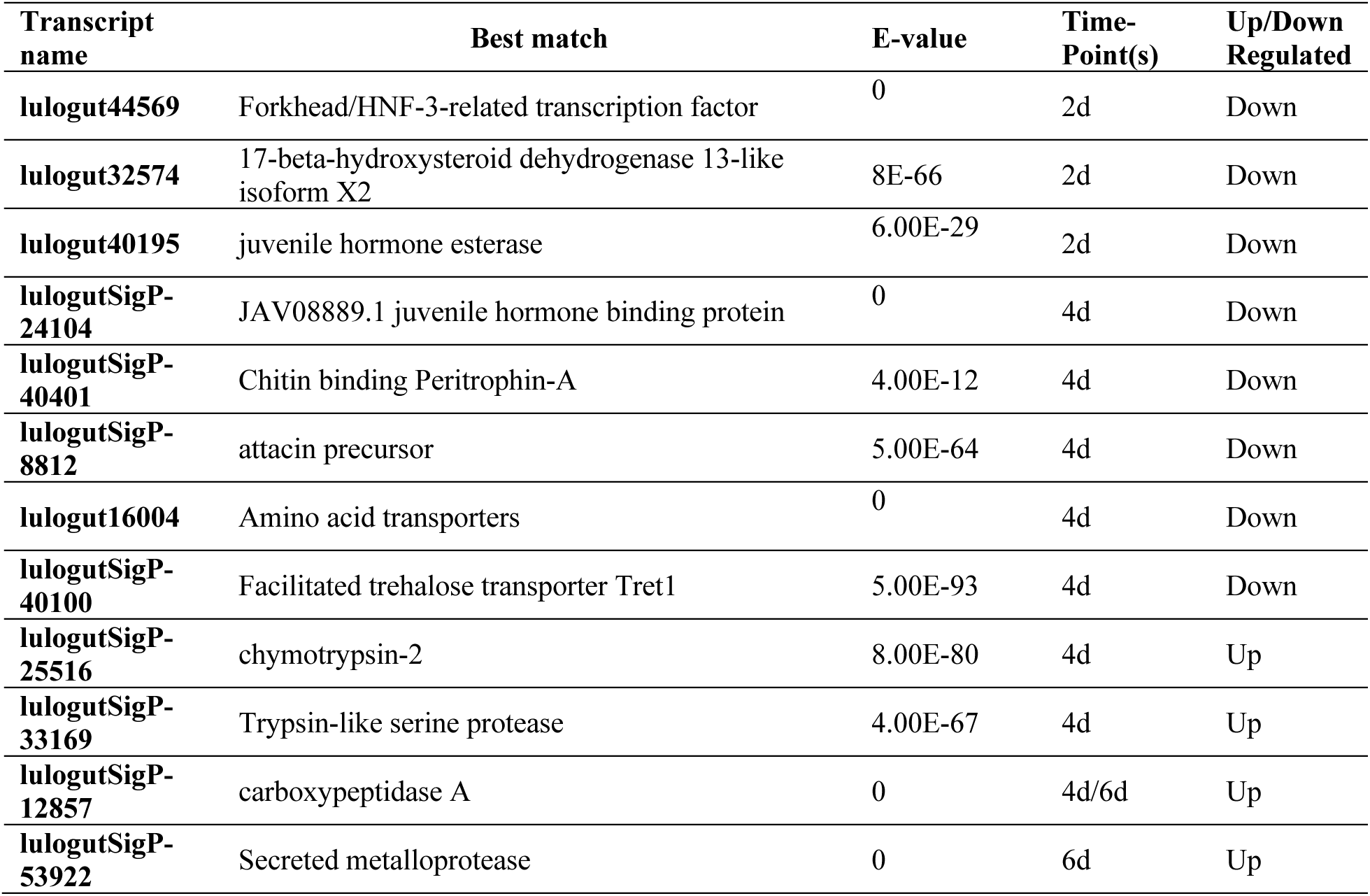
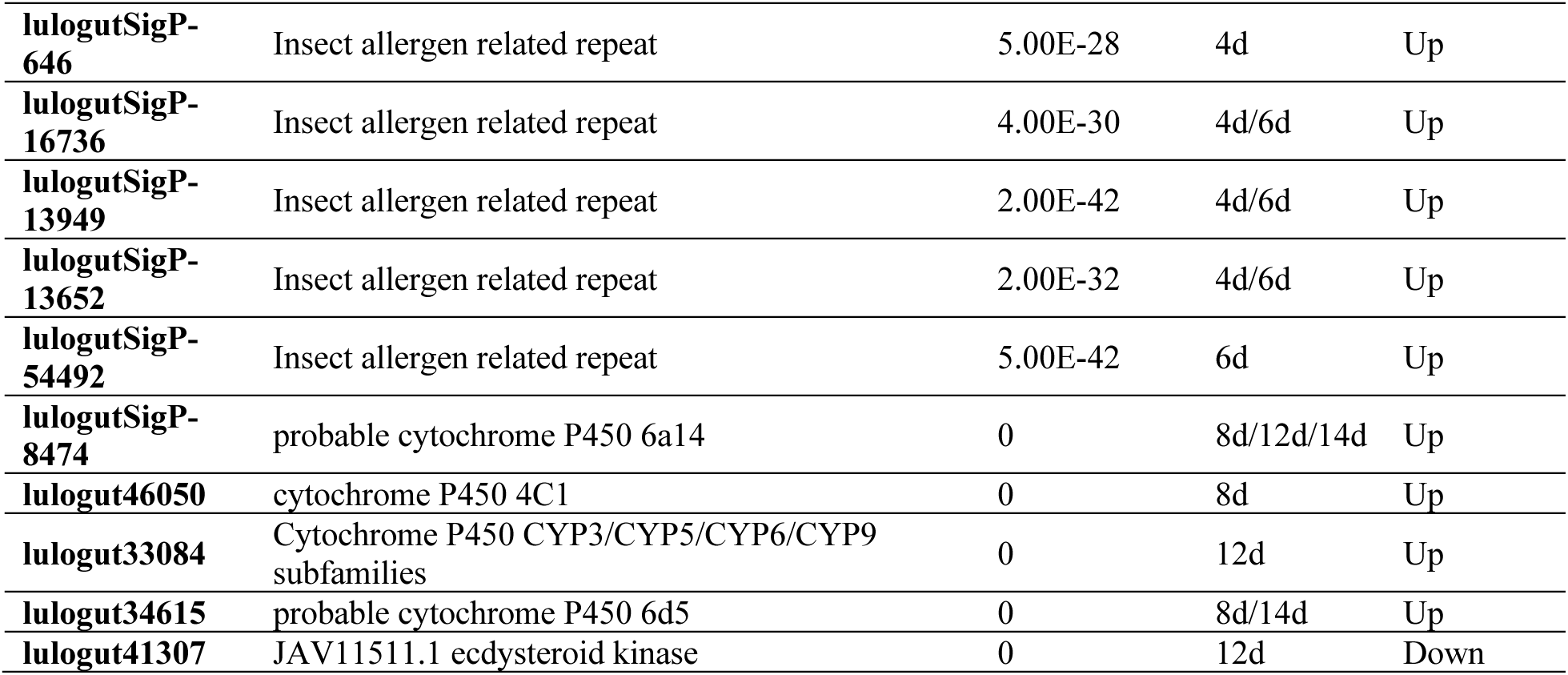
Selected midgut transcripts differentially regulated upon *Leishmania* infection.

| Transcript name | Best match | E-value | Time-Point(s) | Up/Down Regulated |
| --- | --- | --- | --- | --- |
| **lulogut44569** | Forkhead/HNF-3-related transcription factor | 0 | 2d | Down |
| **lulogut32574** | 17-beta-hydroxysteroid dehydrogenase 13-like isoform X2 | 8E-66 | 2d | Down |
| **lulogut40195** | juvenile hormone esterase | 6.00E-29 | 2d | Down |
| **lulogutSigP-24104** | JAV08889.1 juvenile hormone binding protein | 0 | 4d | Down |
| **lulogutSigP-40401** | Chitin binding Peritrophin-A | 4.00E-12 | 4d | Down |
| **lulogutSigP-8812** | attacin precursor | 5.00E-64 | 4d | Down |
| **lulogut16004** | Amino acid transporters | 0 | 4d | Down |
| **lulogutSigP-40100** | Facilitated trehalose transporter Tret1 | 5.00E-93 | 4d | Down |
| **lulogutSigP-25516** | chymotrypsin-2 | 8.00E-80 | 4d | Up |
| **lulogutSigP-33169** | Trypsin-like serine protease | 4.00E-67 | 4d | Up |
| **lulogutSigP-12857** | carboxypeptidase A | 0 | 4d/6d | Up |
| **lulogutSigP-53922** | Secreted metalloprotease | 0 | 6d | Up |
| **lulogutSigP-646** | Insect allergen related repeat | 5.00E-28 | 4d | Up |
| **lulogutSigP-16736** | Insect allergen related repeat | 4.00E-30 | 4d/6d | Up |
| **lulogutSigP-13949** | Insect allergen related repeat | 2.00E-42 | 4d/6d | Up |
| **lulogutSigP-13652** | Insect allergen related repeat | 2.00E-32 | 4d/6d | Up |
| **lulogutSigP-54492** | Insect allergen related repeat | 5.00E-42 | 6d | Up |
| **lulogutSigP-8474** | probable cytochrome P450 6a14 | 0 | 8d/12d/14d | Up |
| **lulogut46050** | cytochrome P450 4C1 | 0 | 8d | Up |
| **lulogut33084** | Cytochrome P450 CYP3/CYP5/CYP6/CYP9 subfamilies | 0 | 12d | Up |
| **lulogut34615** | probable cytochrome P450 6d5 | 0 | 8d/14d | Up |
| **lulogut41307** | JAV11511.1 ecdysteroid kinase | 0 | 12d | Down |

The presence of *Leishmania* in the midgut led to more genes being down-regulated at d2 and up-regulated at later time points, except on 12d (Fig. 3B and Additional File 8: Table S5 and Additional File 9: Fig. S4). On 1d, 2d, and 4d, early time points, 1, 11, and 30 genes were down-regulated (Fig. 3B and Table 2 and Additional File 8: Table S5 and Additional File 9: Fig. S4) whereas 1, 3, and 23 genes were up-regulated (Fig. 3B and Table 3 and Additional File 8: Table S5 and Additional File 9: Fig. S4), respectively. On 6d and 8d, on the other hand, 20 and 15 genes were up-regulated, yet none were down-regulated (Fig. 3B and Additional File 9: Fig. S4). Infected midguts on day 12 displayed 13 up-regulated genes compared to 18 down-regulated ones (Fig. 3B and Additional File 9: Fig. S4). The 14d time point exhibited more up-regulated (12 genes) than down-regulated (1 gene) genes in infected over uninfected midguts (Fig. 3B and Additional File 9: Fig. S4).

**Table 2.**
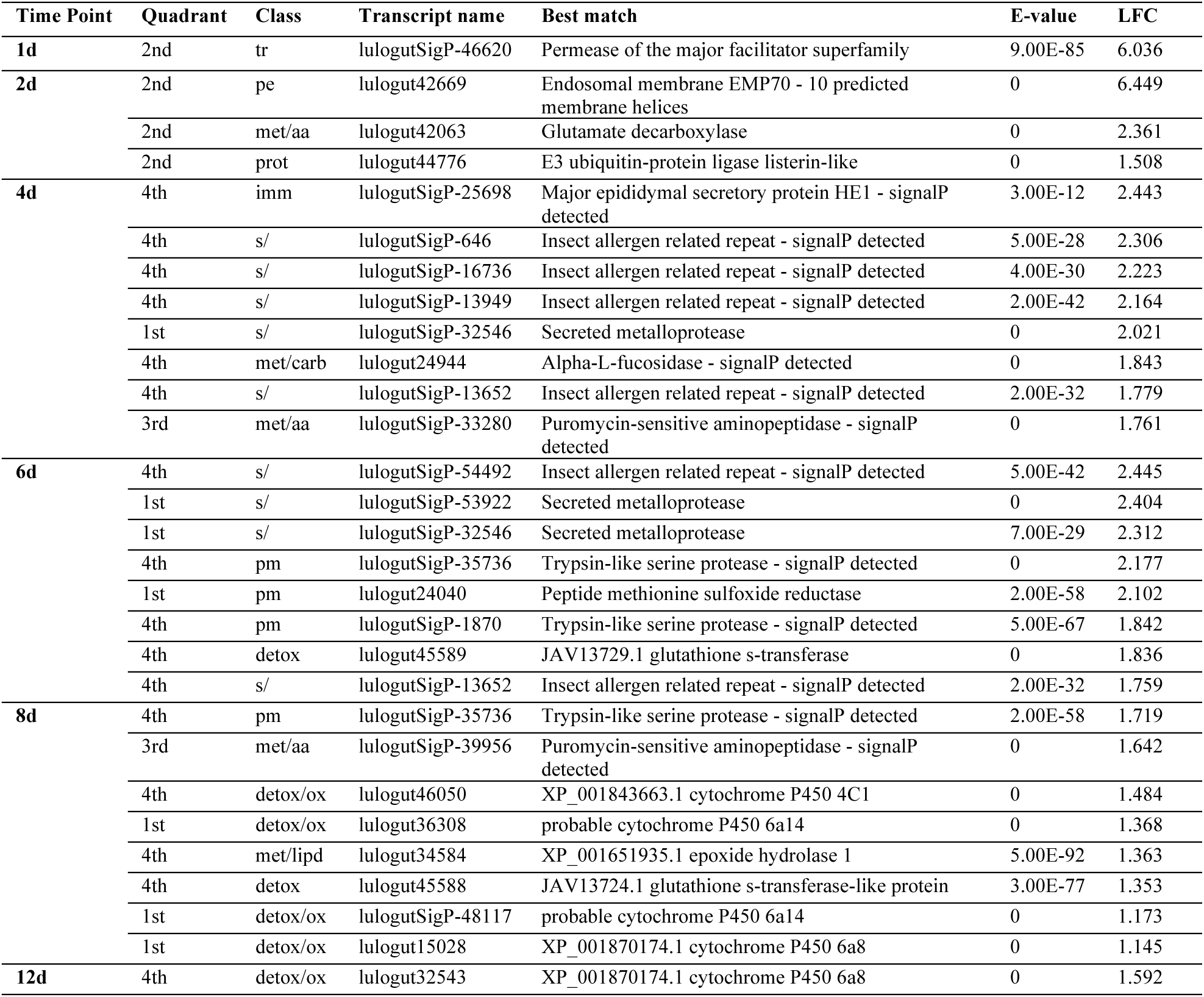
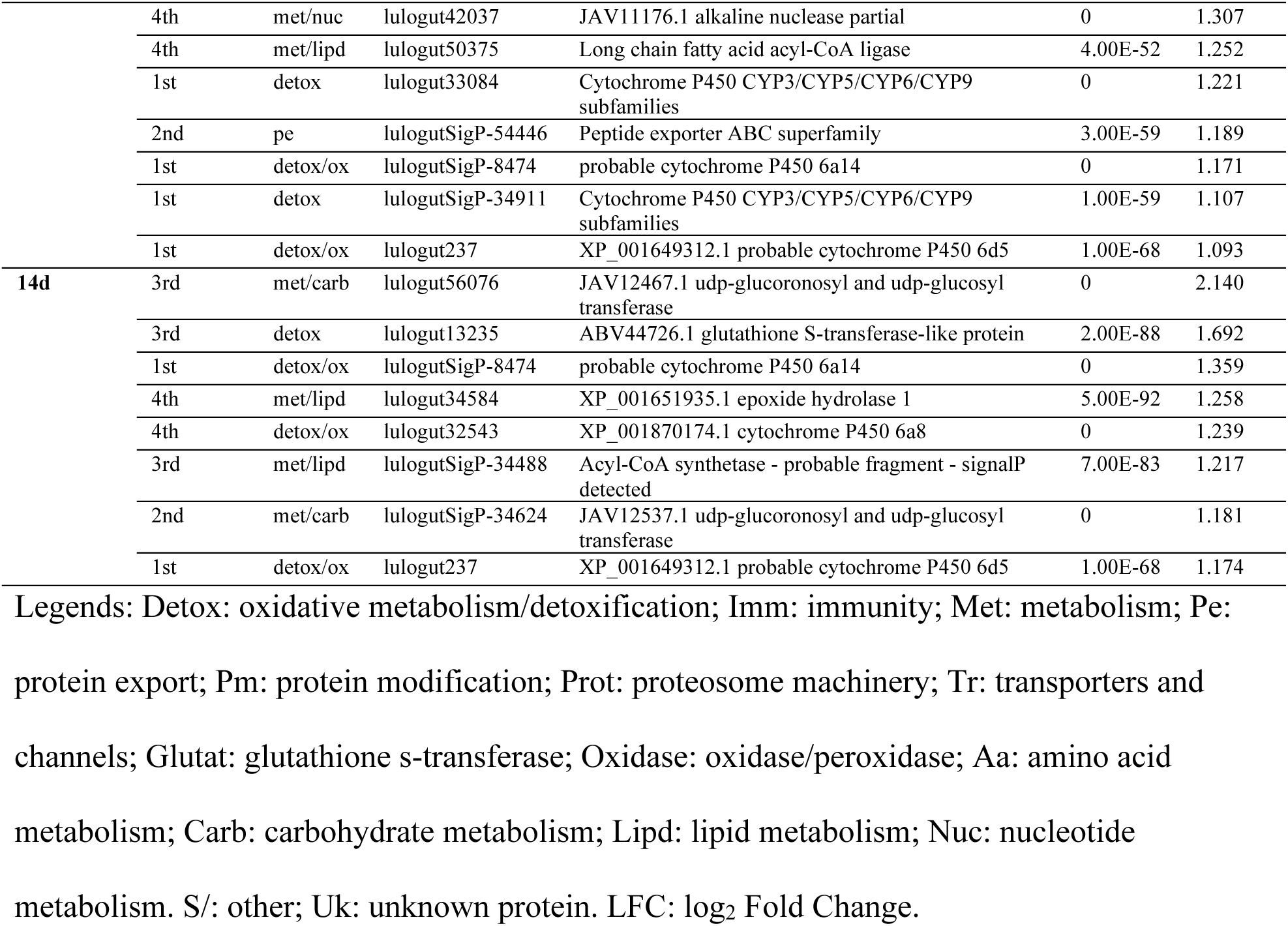
Top eight up-regulated midgut transcripts upon *Leishmania* infection per time point.

**Table 3.** Top five down-regulated midgut transcripts upon *Leishmania* infection per time point.

| Time Point | Quadrant | Class | Transcript name | Best match | E-value | LFC |
| --- | --- | --- | --- | --- | --- | --- |
| 1d | 2nd | nr | lulogut42801 | DNA damage-responsive repressor GIS1/RPH1 jumonji superfamily | 0 | -1.419 |
| 2d | 4th | st | lulogut40195 | NP_523758.3 juvenile hormone esterase isoform A | 6.00E-29 | -1.823 |
|  | 2nd | tr | lulogutSigP-32510 | Permease of the major facilitator superfamily | 0 | -1.9194 |
|  | 2nd | tr | lulogutSigP-46620 | Permease of the major facilitator superfamily | 9.00E-85 | -2.538 |
|  | 2nd | tf | lulogut44569 | Forkhead/HNF-3-related transcription factor | 3.00E-90 | -2.960 |
|  | 3rd | tr | lulogut21743 | JAV05033.1 sodium/potassium-transporting atpase subunit beta-2-like protein | 0 | -2.991 |
|  | 3rd | st | lulogutSigP-22907 | Acetylcholinesterase/Butyrylcholinesterase | 4.00E-54 | -5.397 |
|  | 3rd | s/met/lipd | lulogutSigP-23161 | AAO22149.1 mammalian-like lipase | 0 | -5.688 |
|  | 3rd | uk | lulogutSigP-18032 | Unknown product | NA | -5.917 |
| 4d | 1st | s/uk | lulogutSigP-14897 | hypothetical secreted protein precursor | 1000 | -3.861 |
|  | 3rd | met/lipd | lulogut21836 | JAV11771.1 lipid storage droplet surface-binding protein 1 | 0 | -3.861 |
|  | 3rd | s/met/nuc | lulogutSigP-26492 | JAV11299.1 deoxyribonuclease i partial | 0 | -4.105 |
|  | 2nd | s/protin | lulogutSigP-16416 | BPTI/Kunitz family of serine protease inhibitors | 8.00E-17 | -4.125 |
|  | 2nd | met/carb | lulogut25316 | Hexokinase | 0 | -4.299 |
|  | 2nd | s/ uk | lulogutSigP-16502 | hypothetical conserved secreted protein precursor | NA | -4.926 |
|  | 3rd | s/uk | lulogut36242 | hypothetical secreted protein precursor | 1000 | -5.523 |
|  | 3rd | s/ | lulogutSigP-24104 | JAV08889.1 juvenile hormone binding protein in insects | 0 | -8.423 |
| **12d** | 1st | detox | lulogut19743 | JAV03807.1 metallothionein-2-like protein | 2.00E-34 | -2.909 |
|  | 2nd | storage | lulogut21324 | JAV06440.1 ovotransferrin partial | 0 | -3.778 |
|  | 1st | s/uk | lulogutSigP-16502 | hypothetical conserved secreted protein precursor | NA | -3.893 |
|  | 2nd | s/protinh | lulogutSigP-16416 | BPTI/Kunitz family of serine protease inhibitors - signalP detected | 8.00E-17 | -3.902 |
|  | 2nd | tm | lulogutSigP-15657 | nucleolar and coiled-body phosphoprotein 1 isoform X2 Drosophila ficusphila | 4.00E-21 | -4.086 |
|  | 2nd | pm/protease | lulogut25198 | JAV08757.1 trypsin | 0 | -4.383 |
|  | 2nd | s/ | lulogutSigP-24035 | JAV08413.1 secreted mucin | 0 | -4.536 |
|  | 2nd | met/lipd | lulogut41307 | JAV11511.1 ecdysteroid kinase | 0 | -6.148 |
| **14d** | 2nd | met/nuc | lulogut40330 | Uridylate kinase/adenylate kinase | 4E-59 | -1.292 |

Legends: Detox: oxidative metabolism/detoxificationNr: nuclear regulation; Pm: protein modification; S: secreted protein; St: signal transduction; Storage: storage protein; Tf: transcription factor; Tm: transcription machinery; Tr: transporters and channels; Uk: unknown protein. Met/Carb: carbohydrate metabolism; Met/Lipd: lipid metabolism; Met/Nuc: nucleotide metabolism. S/: other; Protea: protease; Protinh: protease inhibitor. LFC: log<sub>2</sub> Fold Change.

Venn diagrams show that most of genes were differentially expressed at specific time points (Fig. 3C and Additional File 10: Table S6). In the comparisons between early time points (1d through 6d; Fig. 3C, left panel), 1 out of the 2 DE genes on 1d was only modulated at that time point (Fig. 3C, left panel). Similarly, 13 out of the 14 genes, and 43 out of 54 genes, were uniquely DE on 2d and 4d, respectively (Fig. 3C, left panel). Only the 6d DE genes exhibited as many unique as shared with 4d DE genes (10 genes; Fig. 3C, left panel). The comparisons of DE genes between later time points (6d through 14d) showed a greater number of shared DE genes between time points (Fig. 3C, right panel). For instance, only 5 out of 15, and 5 out of 13, DE genes were unique to 8d and 14d, respectively (Fig. 3C, right panel). The 12d midguts, on the other hand, exhibited 26 uniquely expressed genes out 32, the most amongst the late time points (Fig. 2C, right panel).

The expression patterns of all DE genes across time points were assessed through PCA analysis (Fig. 3D and Additional file 11: Table S7). The 113 DE genes were mapped onto a two-dimensional space (expression space), whereby DE genes located close together displayed similar expression profiles through time than those that mapped farther away (Fig 3D). In fact, the DE genes located in the first quadrant of the expression space exhibited about 25-fold greater overall expression levels than those that mapped onto the second and third quadrants (Fig. 3E; Mann Whitney U test, p < 0.0001). Likewise, the DE genes located on the fourth quadrant of the expression space exhibited about 177-fold higher overall expression levels than those that mapped onto the second and third quadrants (Fig. 3E; Mann Whitney U test, p < 0.0001). Looked at through time, the location of the DE genes in different quadrants further highlighted temporal expression differences in both early blood-fed infected midguts and late time point infected midguts (Fig. 3F; Additional file 12: Fig. S5). For example, the DE genes mapped onto the first and second quadrants were either down-regulated at early time points (1d and/or 2d) and up-regulated at later time points (d4 onwards; Fig. 3F and Additional file 12: Fig. S5). On the other hand, the DE genes located on the third and fourth quadrants were up-regulated at 1d and 2d and down-regulated from 4d onwards (Fig. 3F and Additional file 12: Fig. S5). Hence, DE genes located on the first quadrant were expressed at high abundance and lately up-regulated; DE genes mapped onto the second quadrant expressed transcripts at low abundance and were up-regulated at late time points; the third quadrant housed the DE genes expressed at low abundance and up-regulated at early time points; and the DE genes transcribed at high abundance and up-regulated at early time points were localized on the fourth quadrant of the transcriptional space (Fig. 3D).

### Differentially expressed genes at different time points

The up-regulated (Fig. 4A-H and Table 2 and Additional file 13: Table S8) and down-regulated (Fig. 5A-F and Table 3 and Additional file 14: Table S9) DE genes at each time point were plotted onto the transcriptional space in order to assess whether or not the expression of the genes modulated by *Leishmania* across time points followed a specific or a random expression pattern by mapping onto specific quadrants or randomly. All the 113 DE genes were distributed throughout the four quadrants in different proportions: 20%, 37%, 22%, and 21% of the DE genes mapped onto the first through fourth quadrants, respectively (Fig. 4A and Table 2). The up-regulated genes on 1d and 2d were mostly located in the second and third quadrants, which housed genes transcribed at low abundance (Fig. 4B-C and Table 2). However, the reduced gene counts at 1d and 2d precludes statistical comparisons. On the other hand, the DE genes at 4d through 8d followed specific expression patterns (Chi-square test, p < 0.01; Fig. 4D-F). At such time points, 74% (4d), 85% (6d), 87% (8d) of genes up-regulated by *Leishmania* infection mapped onto either the first or fourth quadrant, which housed genes transcribed at high abundance (Fig. 4D-F and Table 2). Although not statistically significant, mapping at 12d and 14d followed a similar pattern where 85% (12d; Fig. 4G and Table 2) and 58% (14d; Fig. 4H and Table 2) of the genes mapped onto either the first or fourth quadrant. However, the proportion of up-regulated genes that mapped on such quadrants gradually changed through time, with more genes mapping onto the fourth quadrant at earlier time points to more genes mapping onto the first quadrant at later time points (Figs 4F-G and Table 2). For instance, 61% and 65% of the up-regulated genes on 4d and 6d mapped onto the fourth quadrant whereas only 13% and 20% of such genes were located on the first quadrant, respectively (Fig. 4D-E and Table 2). In contrast, on 8d, 47% of the *Leishmania* up-regulated genes were located in the first quadrant whereas 40% of such genes were mapped onto the fourth quadrant (Fig. 4F and Table 2). Thereby, most of the DE midgut genes up-regulated by *Leishmania* infection encompassed highly expressed genes, yet the up-regulated genes were more predominant at early time points (4d and 6d) and the late expressed genes were more predominant at late time points (8d to 14d; Fig. 4D-H and Table 2). Interestingly, most of the midgut genes DE by *Leishmania* infection were up-regulated by up to 32-fold (LFC < 5; Fig. 4A-H and Table 2).

**Figure 4.**
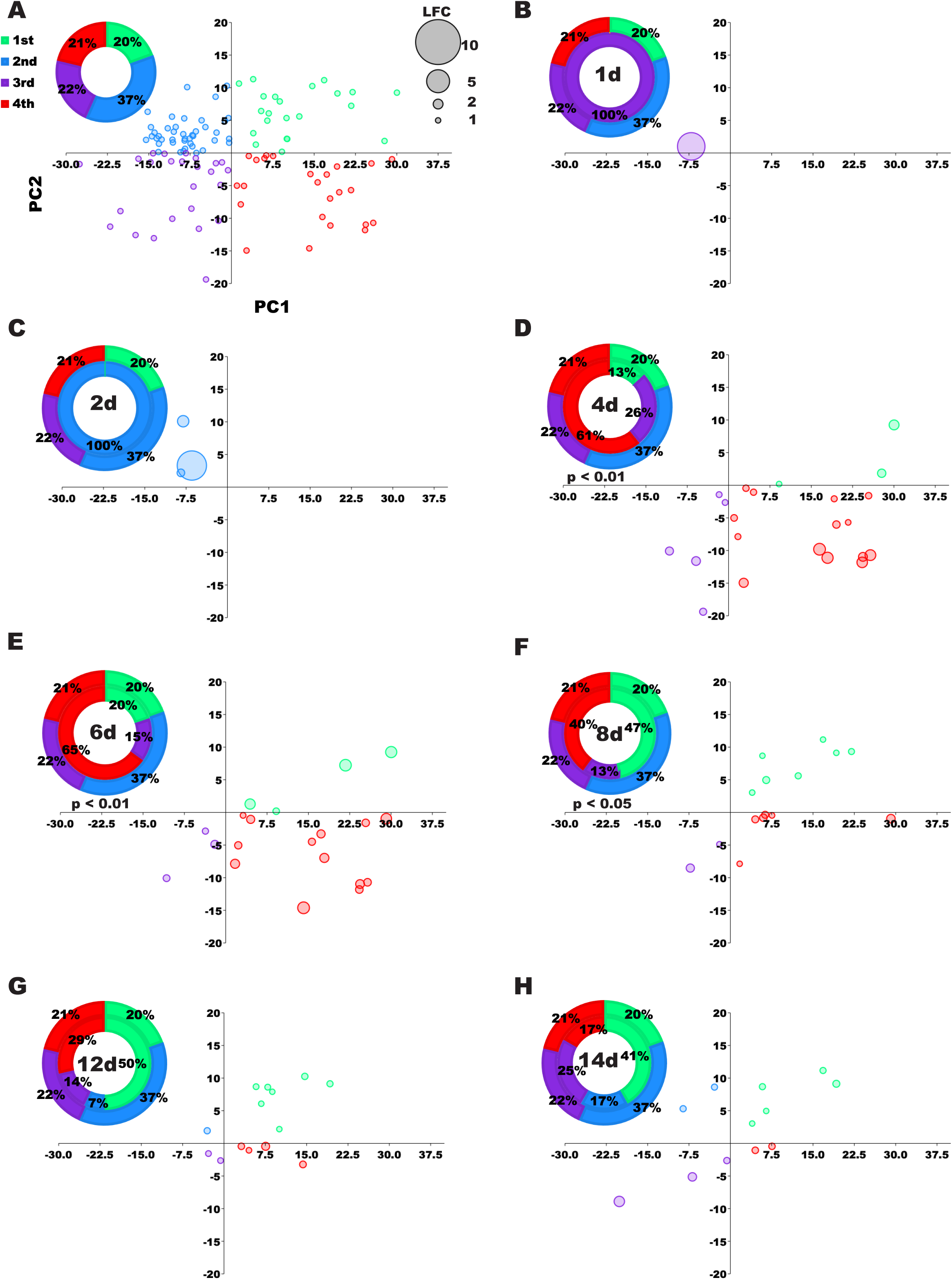
Up-regulated transcripts in *Leishmania*-infected midguts at each time point mapped onto the expression space. **A**. Bubble plot depicts all the DE transcripts mapped onto the expression space. Doughnut chart shows the proportion of transcripts in each quadrant. Inset on the right depicts the scale for the LFC of each up-regulated transcript represented by the diameter of each bubble. **B-H**. Bubble plots mapping the up-regulated transcripts in the expression space for each of the seven time points. The doughnut chart in each graph shows the proportion of up-regulated genes per quadrant (inner circle) and the proportion of all DE genes per quadrant (outer circle), as in A. Differences were statistically significant at p < 0.05 (Chi-square test). DE was considered significant for transcripts displaying LFC higher than 1 and FDR q-value lower than 0.05.

**Figure 5.**
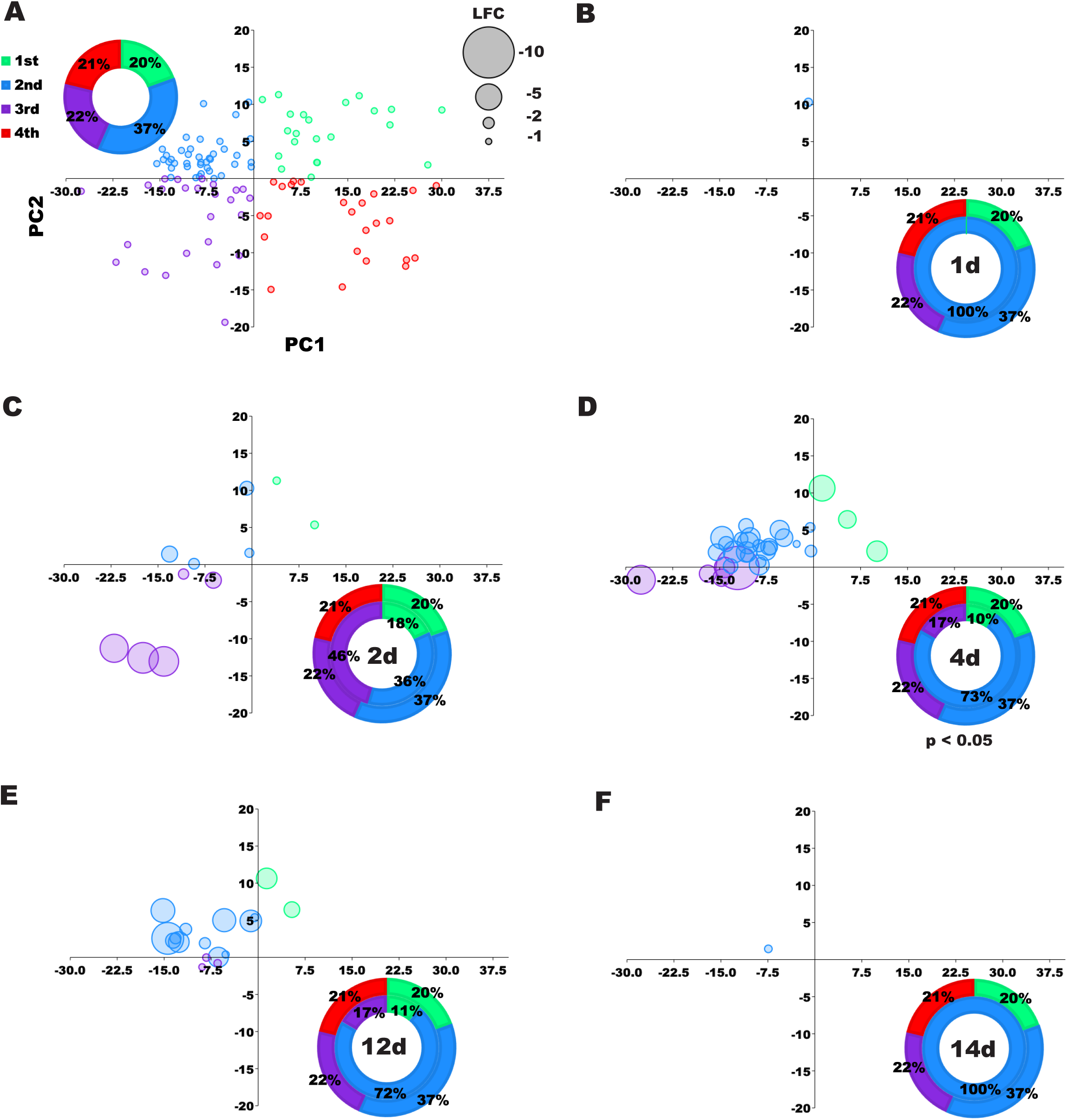
*Leishmania* down-regulated transcripts in each time point mapped on the expression space. **A**. Bubble plot depicts all the DE transcripts mapped onto the expression space. Doughnut chart shows the proportion of transcripts in each quadrant. Inset on the right depicts the scale for the LFC of each down-regulated transcript represented by the diameter of each bubble. **B-F**. Bubble plots mapping the down-regulated transcripts onto the expression space for each of all time points, except days 6 and 8 that were devoid of down-regulated transcripts. The doughnut chart in each graph shows the proportion of down-regulated genes per quadrant (inner circle) and the proportion of all DE genes per quadrant (as in a). Differences were statistically significant at p < 0.05 (Chi-square test). DE was considered significant for transcripts displaying LFC either lower than −1 and FDR q-value lower than 0.05.

Regarding the midgut genes down-regulated by *Leishmania* infection (Fig. 5A-F and Table 3 and Additional file 14: Table S9), none were DE on 6d and 8d (Fig. 5B and Table 3). Contrasting to the midgut up-regulated genes, which exhibited similar expression profiles and were fine-tuned through time (Fig. 5D-F and Table 3), for the most part the midgut down-regulated genes displayed more diverse expression patterns, highlighted by the random distribution of such genes across the transcriptional space on 1d and 2d (Fig. 5B-C; Table 3), and 12d and 14d (Fig. 5 E-F; Table 3). On the other hand, the 4d midguts displayed most of the down-regulated genes on the second quadrant (73%, p < 0.05; Fig. 5D and Table 3), belonging to the group transcribed at low abundance and up-regulated late in infection (Fig. 5D and Table 3). In addition, many of the genes were down-regulated in *Leishmania*-infected midguts by more than 32-fold (LFC > −5; Fig. 5A-F and Table 3).

### Functional profiles of the differentially expressed genes at different time points

Although the midgut genes up- and down-regulated by *Leishmania* infection exhibited different expression patterns across time points (Figs. 4 and 5), such DE genes belonged to the same functional groups for the most part (Fig. 6 and Tables 2 and 3 and Additional file 15: Table S10). Regarding the up-regulated genes, 28%, 38%, and 18% belonged to the detoxification (detox), metabolism (met), and secreted (s) protein molecular functions, respectively (Fig. 6A and Table 2). In fact, the enrichment of such molecular functions amongst the up-regulated genes was consistent through time (Fig. 6B and Table 2): between 2d through 14d for the metabolism function; and between 8d and 14d for the detoxification function. For the secreted protein category, the enrichment of up-regulated genes was more restricted to 4d and 6d (Fig. 6B and Table 2). At earlier time points (1d and 2d), the few up-regulated genes perform different functions ranging from transporter channels (tr, 1d) to proteosome machinery (prot, 2d; Fig. 6B and Table 2). Regarding midgut genes down-regulated by the *Leishmania* infection, 34% of these genes belonged to the metabolism (22%) and secreted protein (12%) functional groups (Fig. 6C and Table 3). Both categories were consistently enriched on 4d, 12d, and 14d (Fig. 6D and Table 3). At earlier time points (1d and 2d), transporter channels (tr, 1d and 2d) and signaling transduction (st, 2d) were the most enriched molecular functions amongst the down-regulated genes (Fig. 6D and Table 3). All the molecular functions identified across time points were matched by analogous GO terms (Additional file 16: Table S11 and Additional file 16: Table S12).

**Figure 6.**
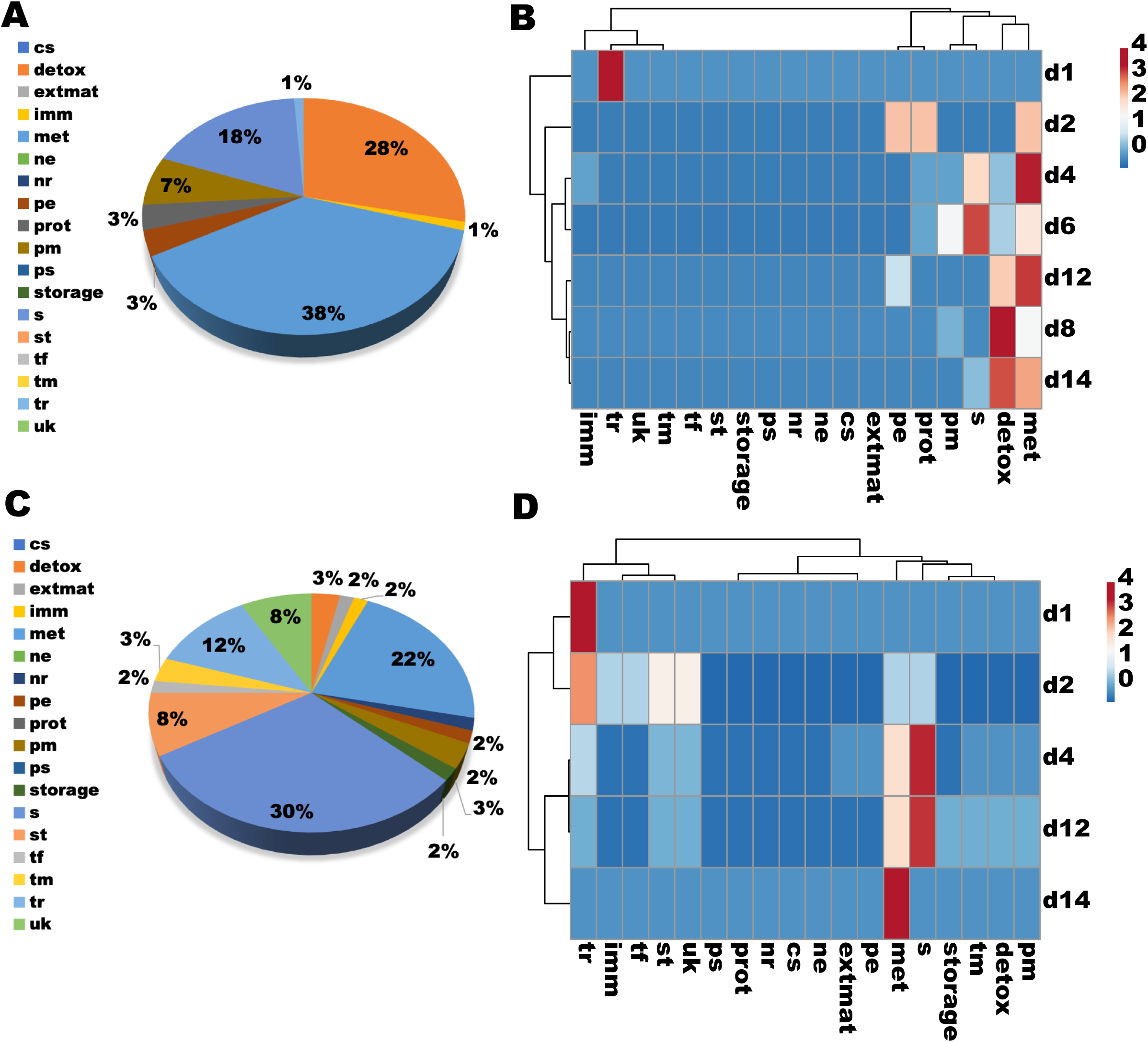
DE transcripts sorted by molecular functions. **A and C.** Pie charts displaying the proportion of midgut DE genes up-regulated (A) and down-regulated (C) by *Leishmania* infection, belonging to different functional groups. **B and D.** Heatmaps and cluster analyses depicting differences in the number of DE genes up-regulated (B) and down-regulated (D) by *Leishmania* infection belonging to different groups of molecular function. Pie chart legends: Cs: cytoskeleton; Detox: oxidative metabolism/detoxification; Extmat: extracellular matrix; Imm: immunity; Met: metabolism; Ne: nuclear export; Nr: nuclear regulation; Pe: protein export; Pm: protein modification; Prot: proteosome machinery; Ps: protein synthesis machinery; S: secreted protein; St: signal transduction; Storage: storage protein; Te: transposable element; Tf: transcription factor; Tm: transcription machinery; Tr: transporters and channels; Uk: unknown protein. The heatmaps are color-coded according to the legends on the right. DE was considered significant for transcripts displaying LFC either lower than −1 or higher than 1 and FDR q-value lower than 0.05.

In order to investigate in-depth the functional profiles of the DE genes, we broke down the most predominant functional classes into subclasses. For the midgut DE genes belonging to the detoxification molecular function (detox), the cytochrome P450 gene family encompassed 76% of the up-regulated genes (Fig. 7A and Table 2 and Additional file 15: Table S10). Such genes were consistently up-regulated between 6d and 14d (Fig. 7B and Table 2). In contrast, the down-regulated genes belonging to the detoxification molecular function were enriched in metallothioneins (4d and 12d, thio; Fig. 7C and D and Table 3). As far as the DE midgut genes belonging to the metabolism function, 55% of the up-regulated genes were related to the metabolism of lipids (lipd; Fig. 8A and Table 2) which was consistently the most predominant between 6d and 14d (Fig. 8B). Among the down-regulated genes performing metabolic functions (Fig. 8C-D and Table 3), most (31%) participated in the metabolism of lipids (lipd) at early time points (2d and 4d) or nucleotides (nuc) on 12d, a later time point (14d, Fig. 8C-D and Table 3). Regarding the DE midgut genes encompassing the secreted proteins (Fig. 9), 50% of those up-regulated belonged to the ‘other category’ (s, multiple protein functions) that was enriched in transcripts of the insect allergen proteins (Fig. 9A; Table 2 and Table S9), also known as microvilli proteins. Although the insect allergens, along with the mucins, and to a lesser extent metalloproteases (metal), were more predominant on 4d and 6d (Fig. 9B and Table 2), up-regulated transcripts encoding proteins of unknown function were enriched at 14d, a later time point (Fig. 9B and Table 2). Among the down-regulated transcripts encoding secreted proteins, 44% belonged to the unknown function (31%, uk) and “other” (17%, s) categories (Fig. 9C and Table 3). The “other” category (s) was consistently down-regulated on 4d and 12d (Fig. 9D and Table 3) and was enriched in transcripts encoding juvenile hormone (JH) binding proteins as well as attacin (Table 3). Transcripts of secreted proteins related to the digestion of lipids (met-li) were down-regulated on 2d (Fig. 9D and Table 3).

**Figure 7.**
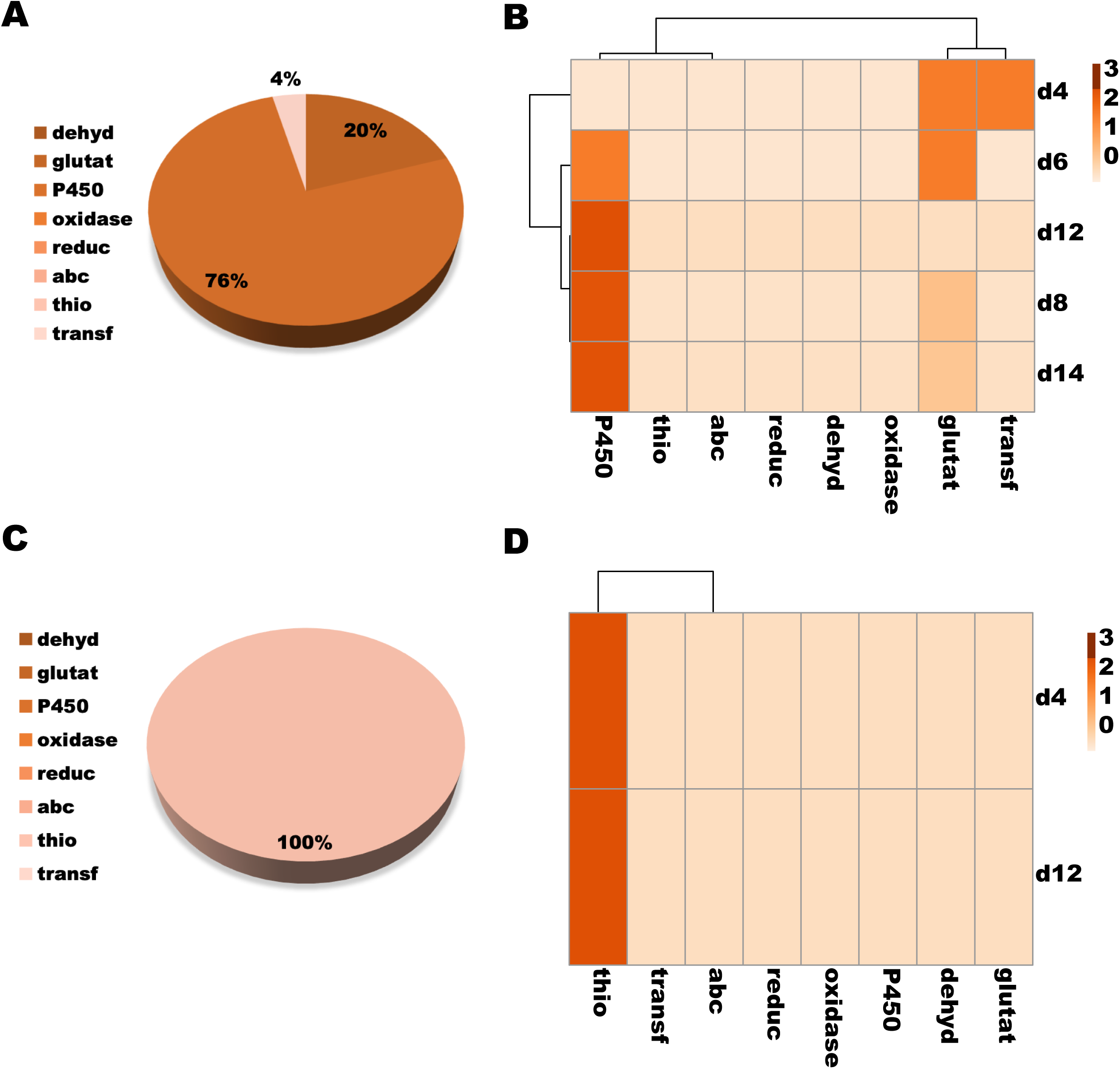
DE transcripts belonging to the molecular function “oxidative metabolism/detoxification” across time points. **A and C.** Pie charts displaying the proportion of midgut DE genes up-regulated (A) and down-regulated (C) by *Leishmania* infection, belonging to the different sorts of oxidative metabolism/detoxification molecular function. **B and D.** Heatmaps and cluster analyses depicting differences in the number of DE genes up-regulated (B) and down-regulated (D) by *Leishmania* infection, belonging to different sorts of oxidative metabolism/detoxification molecular function. Pie chart legends: Dehyd: dehydrogenase; Glutat: glutathione s-transferase; P450: cytochrome P450; Oxidase: oxidase/peroxidase; Reduc: reductase; Abc: Transporter ABC superfamily; Thio: thioredoxin binding protein; Transf: sulfotransferase. The heatmaps are color-coded according to the legends on the right. DE was considered significant for transcripts displaying LFC either lower than −1 or higher than 1 and FDR q-value lower than 0.05.

**Figure 8.**
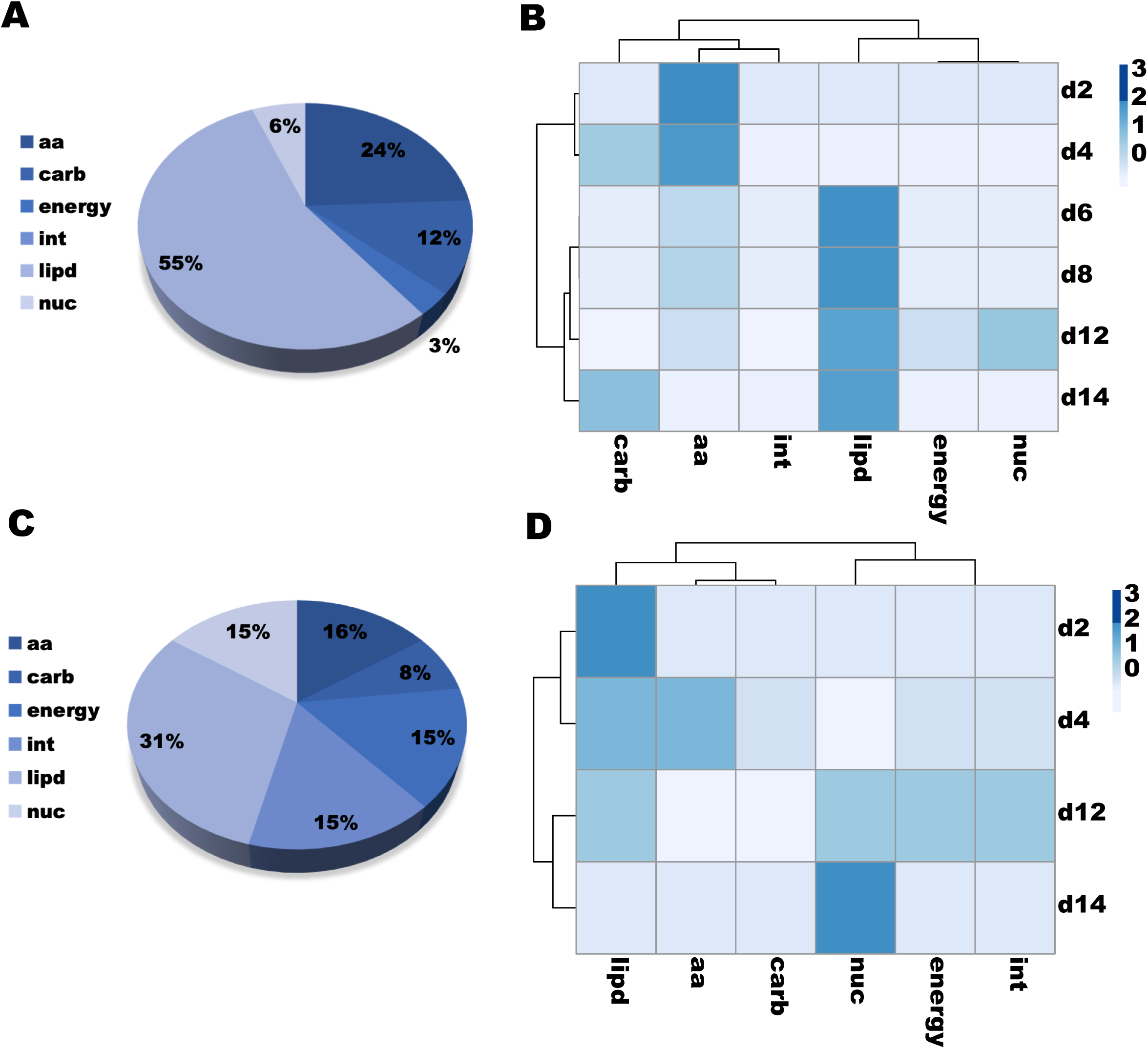
DE transcripts belonging to the molecular function “metabolism” across time points. **A and C.** Pie charts displaying the proportion of midgut DE genes up-regulated (A) and down-regulated (C) by *Leishmania* infection, belonging to the different sorts of metabolism molecular function. **B and D.** Heatmaps and cluster analyses depicting differences in the number of DE genes up-regulated (B) and down-regulated (D) by *Leishmania* infection belonging to different sorts of metabolism molecular function, respectively. Pie chart legends: Aa: amino acid metabolism; Carb: carbohydrate metabolism; Energy: energy production; Int: intermediate metabolism; Lipd: lipid metabolism; Nuc: nucleotide metabolism. The heatmaps are color-coded according to the legends on the right. DE was considered significant for transcripts displaying LFC either lower than −1 or higher than 1 and FDR q-value lower than 0.05.

**Figure 9.**
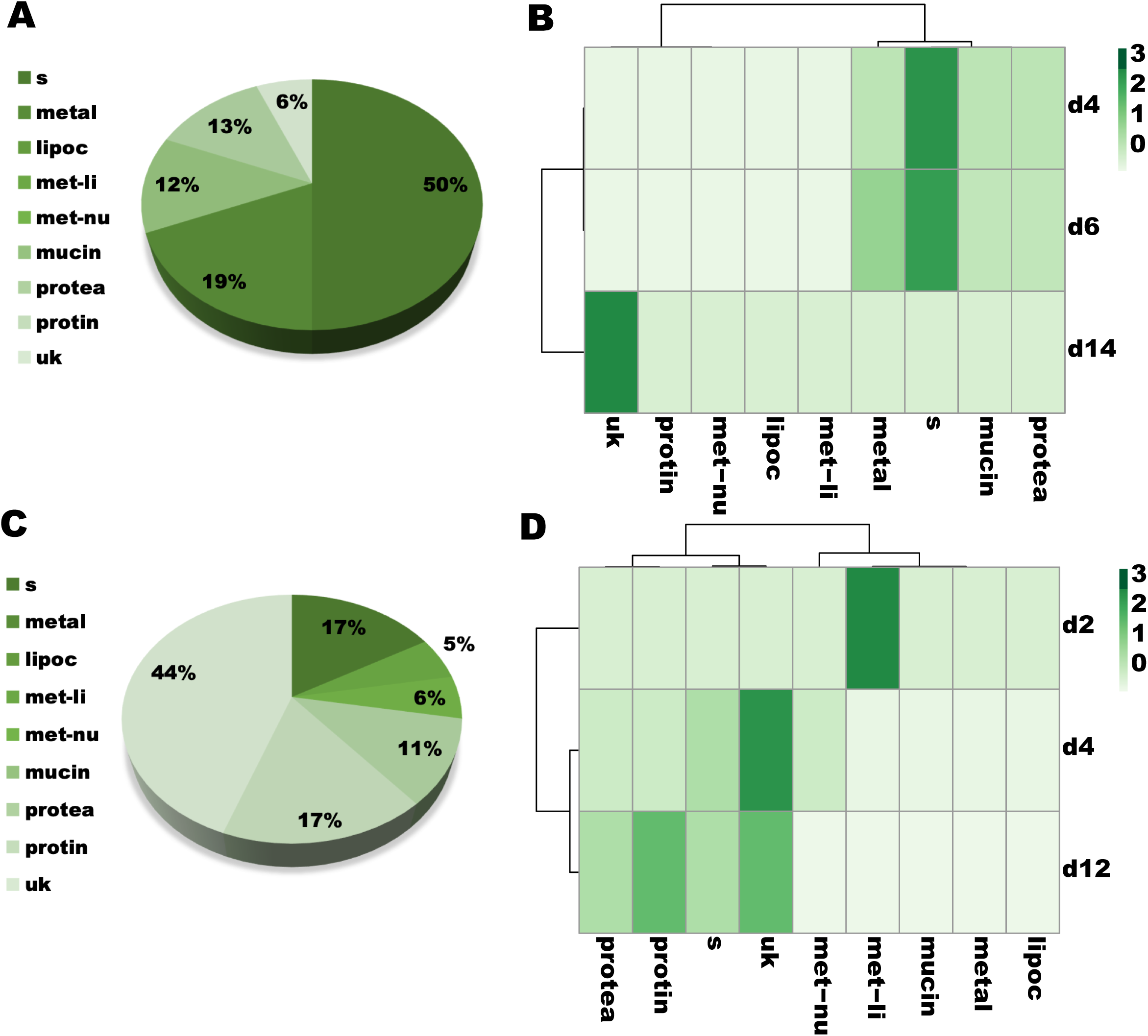
DE transcripts belonging to the molecular function “secreted protein” across time points. **A and C.** Pie charts displaying the proportion of *Leishmania* up-regulated (A) and down-regulated (C) transcripts belonging to the different sorts of secreted protein molecular function. Legends: S: other; Metal: metalloprotease; Lipoc: lipocalin; Met-li: lipase; Met-nu: nuclease; Mucin; Protea: protease; Protin: protease inhibitor; Uk: unknown protein. **B and D.** Heatmaps and cluster analyses depicting differences in the number of DE genes up-regulated (B) and down-regulated (D) belonging to different sorts of secreted protein molecular function. The heatmaps are color-coded according to the legends on the right. DE was considered significant for transcripts displaying LFC either lower than −1 or higher than 1 and FDR q-value lower than 0.05.

## Discussion

In this work, we have carried out a broad RNA-Seq investigation to assess the effects of *Leishmania* infection in sand fly midgut gene expression. As no sand fly genome is available at the standards to be used as a reference for read mapping, all the reads obtained were assembled de novo into 13,841 putative transcripts. Such transcripts were then used as a reference for gene expression quantification and comparisons between infected and uninfected samples. Out of seven time points, only about 1% of the genes were differentially expressed (113 genes) by *Leishmania* infection, highlighting the extent of the adaptation of *Le. infantum* to its natural vector, the sand fly *Lu. longipalpis*.

Multiple midgut genes displaying differential expression upon Leishmania infection in cDNA libraries of *Le. infantum*-infected *Lu. longipalpis* midguts [19] were also differentially expressed in our RNA-Seq libraries. For instance, all four insect allergen proteins (microvilli proteins), multiple digestive enzymes (proteases and peptidases), an astacin-metalloprotease, as well as a peritrophic matrix protein were differentially regulated by Leishmania infection in both studies [19].

The limited influence of *Leishmania* in midgut gene expression as observed in this study was further investigated by PC analysis. As indicate by PC1, most of the variance (77%) in the transcriptional levels across midgut samples was caused by the presence (or lack of) blood in the midguts, sorting out the early (d1 and d2; blood engorged) from late (d4 onwards; blood passed) time points. Even though, PC2 (6.4%) and PC3 (4.1%) exhibited similar levels of variance, PC3 accounted for most of the variance sorting infected from uninfected midguts, and likely represents the differential expression of the 113 genes modulated by *Leishmania* infection. These findings also suggest that other factors not controlled for by the experimental design accounted for the variance observed in PC2. Along these lines, it is noteworthy that Leishmania infection in sand fly midguts also modify the microbiota composition [17], which may also have affected gene expression in the midgut samples.

It is worth noting that multiple genes DE upon *Leishmania* infection were unique to a particular time point, being more pronounced in early time points. This phenomenon may be explained by the enrichment of different *Leishmania* stages at specific time points. For instance, time points 1d, 2d, 4d, and 6d are enriched in amastigotes and transitional stages, and procyclic, nectomonad, and leptomonad promastigotes, respectively. From 6d onwards, *Leishmania* parasites undergo metacyclogenesis: hence, there is a gradual increase in the proportions of metacyclic compared to leptomonad promastigotes through time, which can explain the overlap of DE genes between midguts on 8d and the other late time points. Surprisingly, we observed a burst of down-regulated DE genes on 12d that was not observed on 14d. At both time points the midgut infection is very similar as far as parasite stage and density, a phenomenon that needs to be further investigated.

In order to complete its life cycle in the sand fly midgut, *Leishmania* needs not only to develop and differentiate into the infective metacyclic stage, but also to escape the barriers imposed by the sand fly midgut early in the infection (day 1-5). During this period, *Leishmania* needs to shield itself against the harmful actions of the proteolytic enzymes [9], avoid the immune system [10, 11], escape from the peritrophic matrix [12, 13], and attach to the midgut epithelium [14]. At these early time points, most of the sand fly DE genes were down-regulated by large fold changes. Such sand fly genes are transcribed at high abundance for the most part. On day 4, multiple sand fly genes encoding digestive enzyme as well as a peritrophic matrix protein were down-regulated, pointing to parasite manipulation of the barriers imposed by the sand fly midgut in order to survive. Along the same lines, it is important to highlight that the presence of *Leishmania* in the sand fly midgut leads to the down regulation of genes potentially involved with the control of gene expression. For instance, among the sand fly transcripts down-regulated on day 2 is the transcription factor Forkhead/HNF-3, which is involved with midgut regeneration [22], and nutrient transport and absorption [23]. Accordingly, we have also observed down-regulation of sand fly amino acid and trehalose transporters on 4d after *Leishmania* infection. Transcripts for metallothionein-2-like protein were also down-regulated at the same time point. The expression levels of these proteins are used as a proxy of heavy metals absorption [24]. Hence, their down-regulation in *Leishmania*-infected midguts suggests that these parasites reduce nutrient uptake by the sand fly midgut epithelium. Along the same lines, genes encoding proteins associated with metabolism of hormones, such as the juvenile hormone and ecdysone, were down-regulated on days 4 and 6. Such hormone levels change during blood digestion [25], and relevantly control the expression of sand fly midgut serine proteases [26–28], which are also down-regulated upon *Leishmania* infection on days 4 and 6. Together, these data suggest that the sand fly transcription factor Forkhead/HNF-3 as well as hormone metabolic enzymes might be key targets to control *Leishmania* infection early on.

As the remains of the digested blood is flushed out and the parasites detach from the epithelium [14], the parasites undergo metacyclogenesis from day 6 onwards, migrating to the anterior midgut and differentiating into infective forms [8]. At this late period in the infection, midgut barriers to *Leishmania* development are unknown or negligible. The parasites seem to multiply freely secreting a massive amount of carbohydrates (fPPG) that jams the blood intake and allows the parasites to be regurgitated into the skin [29, 30]. Most of the sand fly DE genes late in infection (day 8 onwards) were up-regulated by narrow fold change differences in response to *Leishmania*. Such genes are transcribed at high abundance for the most part. Most of these genes encode proteins that participate in detoxification of xenobiotics (cytochrome P450) and metabolism of lipids. At these time points, it seems plausible that the massive amount of parasites, reaching 120,000 cells on average on day 14 [21], might be indirectly modulating sand fly gene expression by the release of cell membranes and metabolites from dead parasites and *Leishmania*-derived exosomes [31] throughout metacyclogenesis. Interestingly, the presence of *Leishmania* is undetected by the midgut immune system of the sand fly during this period. This also noted at early time points with the exception of day 4 where the down-regulation of a gene encoding attacin, an antimicrobial peptide [32], was observed. The lack of *Leishmania* detection by the immune system may constitute another adaptation of *Le. infantum* to survive in *Lu. longipalpis* midguts.

## Conclusion

Overall, the presence of *Le. infantum* in the midgut of its natural vector has direct and indirect effects on sand fly midgut gene expression. On one hand, these parasites appear to manipulate gene expression early on to weaken developmental barriers imposed by the midgut. On the other hand, *Leishmania* behaves like a commensal later in the infection, and changes in the sand fly gene expression by the parasites seem to be an indirect consequence of the massive amount of the parasites inside the anterior portion of the midgut.

## Methods

### *Leishmania* parasites, parasite load assessment, sand fly blood feeding and infection, and midgut dissection and storage

Sand fly infection and *Leishmania* counts were performed as described in our companion manuscript [21]. As controls, *Lu. longipalpis* sand flies were also fed on uninfected heparinized dog blood at the same time. After feeding, fully fed females were sorted and given 30% sucrose solution *ad libitum*. Sand flies from both groups were dissected with fine needles and tweezers on a glass slide at days one, two, four, six, eight, twelve, and fourteen after blood feeding on RNAse Free PBS (1X). Forty to sixty midguts were quickly rinsed in fresh RNAse Free PBS (1X) and stored in RNAlater (Ambion), following manufacturer’s recommendation.

### RNA extraction and quality control

Total RNA was extracted using the PureLink RNA Mini Kit (Life Technologies, Carlsbad), following the manufacturer’s recommendations, as described in the companion manuscript [21].

RNA amounts and purification were assessed using a Nanodrop spectrophotometer (Nano Drop Technologies Inc, Wilmingtom; ND-1000), and quality control was further evaluated using a Bioanalyzer (Agilent Technologies Inc, Santa Clara, CA; 2100 Bioanalyzer), using the Agilent RNA 6000 Nano kit (Agilent Technologies) and following the manufacturer’s recommendations. Only one out of the forty-two samples displayed RIN (RNA integrity number) value lower than 7 (Replicate 3 - 14d Pi – RIN 6.7).

### RNA-Seq library preparation and deep sequencing

The RNA-Seq libraries were prepared using the NEBNext® Ultra™ RNA Library Prep Kit for Illumina (New England Biolabs, Ipswick MA), following manufacture’s recommendation, for Single Ended sequencing by HiSeq 2500 (Illumina, San Diego, CA) of 125 nucleotides reads (SE - 125). The RNA-Seq library preparation and sequencing was performed at the NC State University Genomic Science Laboratory.

### Bioinformatic pipeline and de novo assembly

RNA-seq data trimming and mapping were describe elsewhere [21]. *De novo* assembly from high quality reads were a result of both Abyss (kmers from 21 to 91 in 10-fold increments) and Trinity (V2.1.1) assemblers. The combined fasta files were further assembled using an iterative blast and CAP3 pipeline as previously described [33]. Coding sequences were extracted based in the predicted longer open reading frame or the presence of a signal peptide and by similarities to other proteins found in the Refseq invertebrate database from the National Center for Biotechnology Information (NCBI), proteins from Dipterans deposited at NCBI’s Genbank and from SwissProt. Automated annotation of proteins was based on a vocabulary of nearly 350 words found in matches to various databases, including Swissprot, Gene Ontology, KOG, Pfam, Drosophila mRNA transcripts, Virus, and SMART, Refseq-invertebrates and the Diptera subset of the GenBank sequences obtained by querying diptera [organism] and retrieving all protein sequences. Raw reads were deposited on the Sequence Read Archive (SRA) of the National Center for Biotechnology Information (NCBI). This Transcriptome Shotgun Assembly project has been deposited at DDBJ/EMBL/GenBank and will be available when the paper is accepted. Novel Coding sequences and putative protein sequences were submitted to the NCBI from accession numbers and will be available when the paper is accepted.

Raw reads were mapped to the generated dataset using the RNA-Seq by Expectation Maximization (RSEM) vs 1.3.0, Bowtie vs 2-2.2.5 and samtools vs 1.2[34]. Differential expression among timepoints and conditions were analyzed using the R suite by the Bioconductor package DeSeq2 vs 3.8 [35]. Filtering on all mapped gene counts was performed to exclude genes where the sum of counts in all the conditions was inferior to 10 counts. Default parameters were used with DESeq2 including the shrinks log2 fold-change (FC) estimated for each tested comparison [35, 36]. A log_2_ Fold Change and its standard error were generated in addition to a P-value (p-value) and a P-adj (Adjusted p-value) to account for the false discovery rate. Significant associations were considered when a P-adj was smaller than 5% (p <0.05) and log_2_ fold change larger than 0.5 (+/-).

### Data and statistical analyses

Bubble plots and principal component analyses (PCA) were performed using the PAST3 software [37]. For the later, either the log_2_ TPMs or log_2_ fold change (LFC) were used. Statistical analyses were carried out with Prism 7 (GraphPad Software Inc; all the other tests). Venn diagram results were obtained with Venny 2.1 (http://bioinfogp.cnb.csic.es/tools/venny/), and heatmaps/cluster analyses were obtained using the ClustVis tool ([38]; https://biit.cs.ut.ee/clustvis/). Gene heatmap and volcano plots were obtained with the packages gplots and ggplot2 and constructed with the R software.

### nCounter XT gene expression assessment

Gene expression validation was carried out using the nCounter probe-based hybridization assay (NanoString Technologies Inc, Seattle, WA), following the manufacturer’s recommendation. Forty-two sand fly genes were randomly chosen (Additional file 6: Table S3) for probe design and hybridized against 100 ng of each RNA sample, resulting in three biological replications per time point. Raw output data were analyzed using the nSolver software (NanoString Technologies), normalizing the results against the counts for all 42 genes. Only genes detected by the nCounter were considered for comparisons to RNA-Seq data. For a gene to be considered nCounter-detected [39], the average counts for the experimental gene had to be significantly higher than the average counts of eight negative control by Mann Whitney U test (p < 0.05) in at least one of the treatments (infected or uninfected). The expression of the detected genes in each time point was used for expression comparisons with the RNA-Seq expression results for the correspondent genes. For these comparisons, only genes displaying average TPM of at least 1 in one of the treatments were considered. Fold change correlations were determined by plotting the log_2_ ratio of the infected over the uninfected expression values for RNA-Seq (TPMs) and nCounter (normalized counts) and calculating the linear regression coefficient.

## Declarations

### Ethics approval and consent to participate

Not applicable

### Consent for publication

Not applicable

### Availability of data and material

The datasets used and/or analysed during the current study available from the corresponding author on reasonable request.

## Author Contribution

I.V.C.A. and T.D.S. designed and performed the experiments. F.O. supervised bioinformatic analysis. I.V.C.A analyzed the data. C.M. performed sand fly insectary work. J.G.V., S.K. and F.O. were involved in the design, interpretation and supervision of this study. I.V.C.A wrote the first draft of the manuscript. J.G.V., S.K. and F.O edited the manuscript.

## Acknowledgement

We are also thankful to T.R. Wilson and B.G. Bonilla from LMVR, NIAID for sand fly insectary support.

## Abbreviations

Forkhead/HNF-3: Hepatocyte nuclear factor 3/fork head
TPM: Transcripts per million
PBM: Post blood meal
Pi: Post infection
fPPG: Filamentous proteophosphoglycan
DE: Differentially expressed
PCA: Principal component analysis
LFC: Log 2 fold change
ORF: Open reading frame
GO: Gene ontology
SRA: Sequence Read Archive
NCBI: National Center for Biotechnology Information
PBS: Phosphate buffer saline

## Additional Files

Additional file 1:

**Figure S1** Heatmap displaying the expression profiles and cluster analyses of the midgut transcripts across seven time points in uninfected and *Leishmania*-infected samples. The 10,000 most highly expressed transcripts are depicted.

Additional file 2:

**Table S1** Transcriptional and bioinformatics description of the *Lu. longipalpis* midgut transcripts.

Additional file 3:

**Table S2** Summary of the overall percentage of contigs (% of contigs) or abundance (%TPM) for all time points. The distribution of the mapped reads to the functional classification are highlighted.

Additional file 4:

**Figure S2** Pie chart depicting the overall proportion of transcripts belonging to the same molecular function group. Cs: cytoskeleton; Detox: oxidative metabolism/detoxification; Extmat: extracellular matrix; Imm: immunity; Met: metabolism; Ne: nuclear export; Nr: nuclear regulation; Pe: protein export; Pm: protein modification; Prot: proteosome machinery; Ps: protein synthesis machinery; S: secreted protein; St: signal transduction; Storage: storage protein; Te: transposable element; Tf: transcription factor; Tm: transcription machinery; Tr: transporters and channels; Uk: unknown protein.

Additional file 5:

**Table S3** Principal component analysis output for comparisons between average transcriptional expression amongst time points as well as for individual replicates.

Additional file 6:

**Figure S3** Principal component analysis (PCA) describing the position of each replicate for each midgut time point in the expression space. (A) Expression space was generated based on the log2 of TPMs using the 10,000 most expressed transcripts across libraries. The Eigenvalues and % variance for PC1 and PC3 were 5632.97 and 60% and 321.15 and 3.4%, respectively. (B) Expression space between PC1 and PC2. The Eigenvalues and % variance for PC2 were 670.05 and 7.1%, respectively. The color codes labeling each time point were as follow: B. Aqua (1d); C. Royal Blue (2d); D. Sea Green (4d); E. Sandy Brown (6d); F. Saddle Brown (8d); G. Red (12d); and H. Fuchsia (14d).

Additional file 7:

**Table S4** nCounter probes, counts, and expression comparisons with RNA-Seq TPMs.

Additional file 8:

**Table S5** Gene sets displaying differential gene expression at each time point.

Additional file 9:

**Figure S4** Volcano plots depicting the differentially expressed (DE) transcripts at each time point. (A-G). DE transcripts at 1d, 2d, 4d, 6d, 8d, 12d, and 14d, respectively. Only transcripts exhibiting q-values lower than 0.05 are shown. Transcripts displaying fold change greater or lower than 2 (−1 < LFC > 1) are color coded, as follow: Aqua (1d); Royal Blue (2d); Sea Green (4d); Sandy Brown (6d); Saddle Brown (8d); Red (12d); and Fuchsia (14d). LFC scale is color coded in gray (top right). In black, transcripts not significant at −1 < LFC > 1.

Additional file 10:

**Table S6** Genes uniquely differentially expressed at each time point.

Additional file 11:

**Table S7** Gene sets mapping on each quadrant of the PCA map.

Additional file 12:

**Figure S5** Expression analysis per quadrant per time point in infected libraries (Pi). The average TPM for each time point for every DE transcript mapped in each quadrant was plotted. Mean TPM as shapes and SEM bars are depicted.

Additional file 13:

**Table S8** Sets of up-regulated genes mapping on each quadrant of the PCA map at each time point.

Additional file 14:

**Table S9** Sets of down-regulated genes mapping on each quadrant of the PCA map at each time point.

Additional file 15:

**Table S10** Functional analyses of differentially expressed genes.

Additional file 16:

**Table S11** Gene Ontology (GO) enrichment for the up-regulated genes at each time point.

Additional file 17:

**Table S12** Gene Ontology (GO) enrichment for the down-regulated genes at each time point.

