## Supplementary figures and images for "*Leishmania* infection induces a limited differential gene expression in the sand fly midgut"

### Fig S1

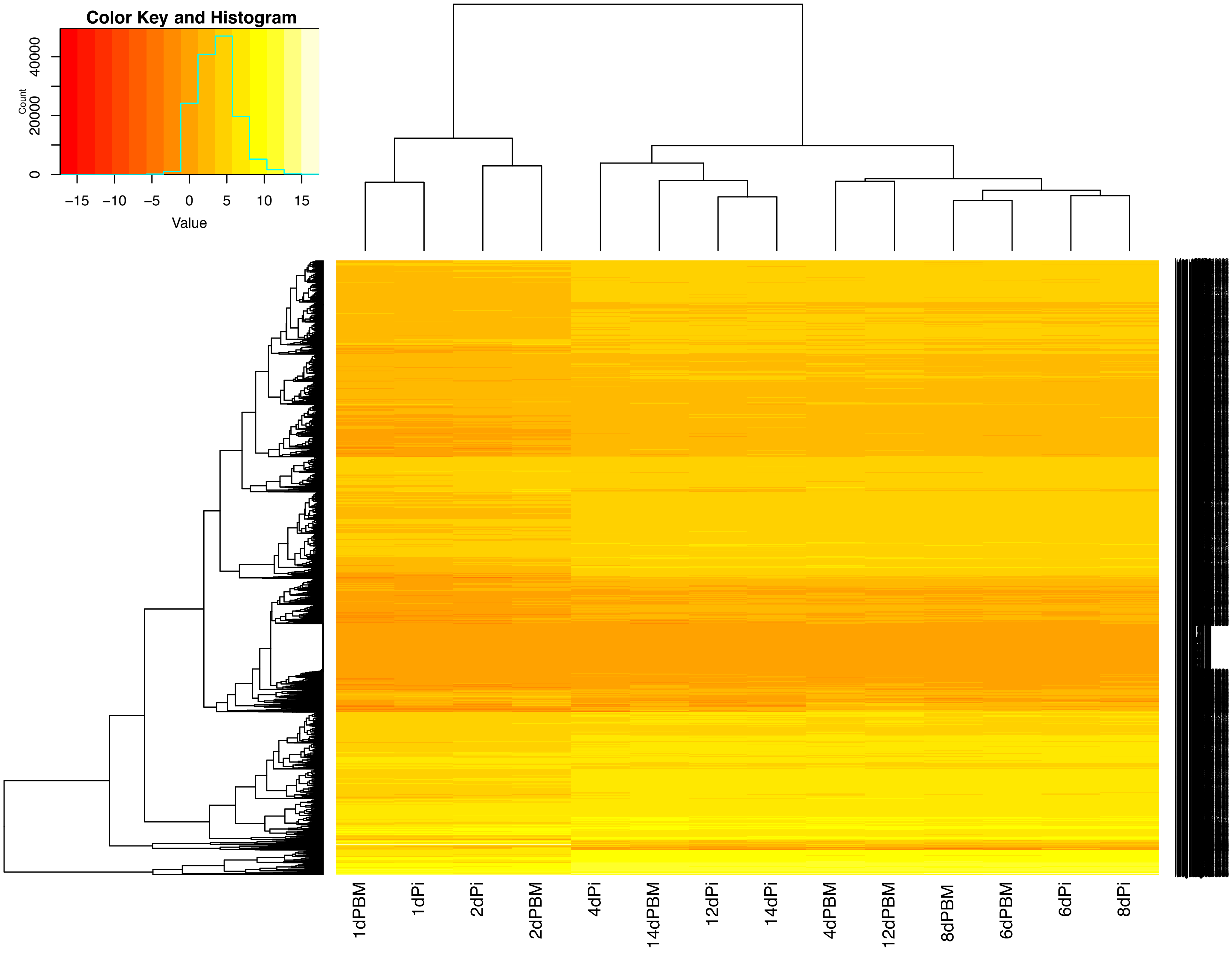

### Fig S2

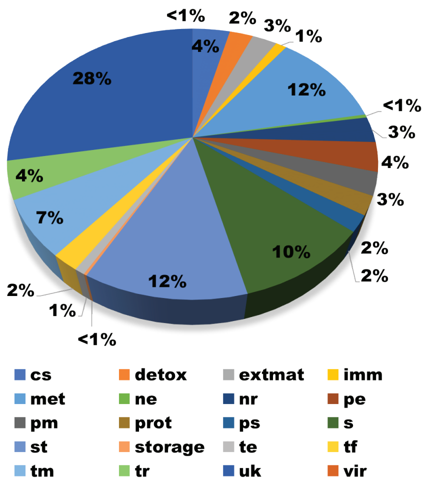

### Fig S3

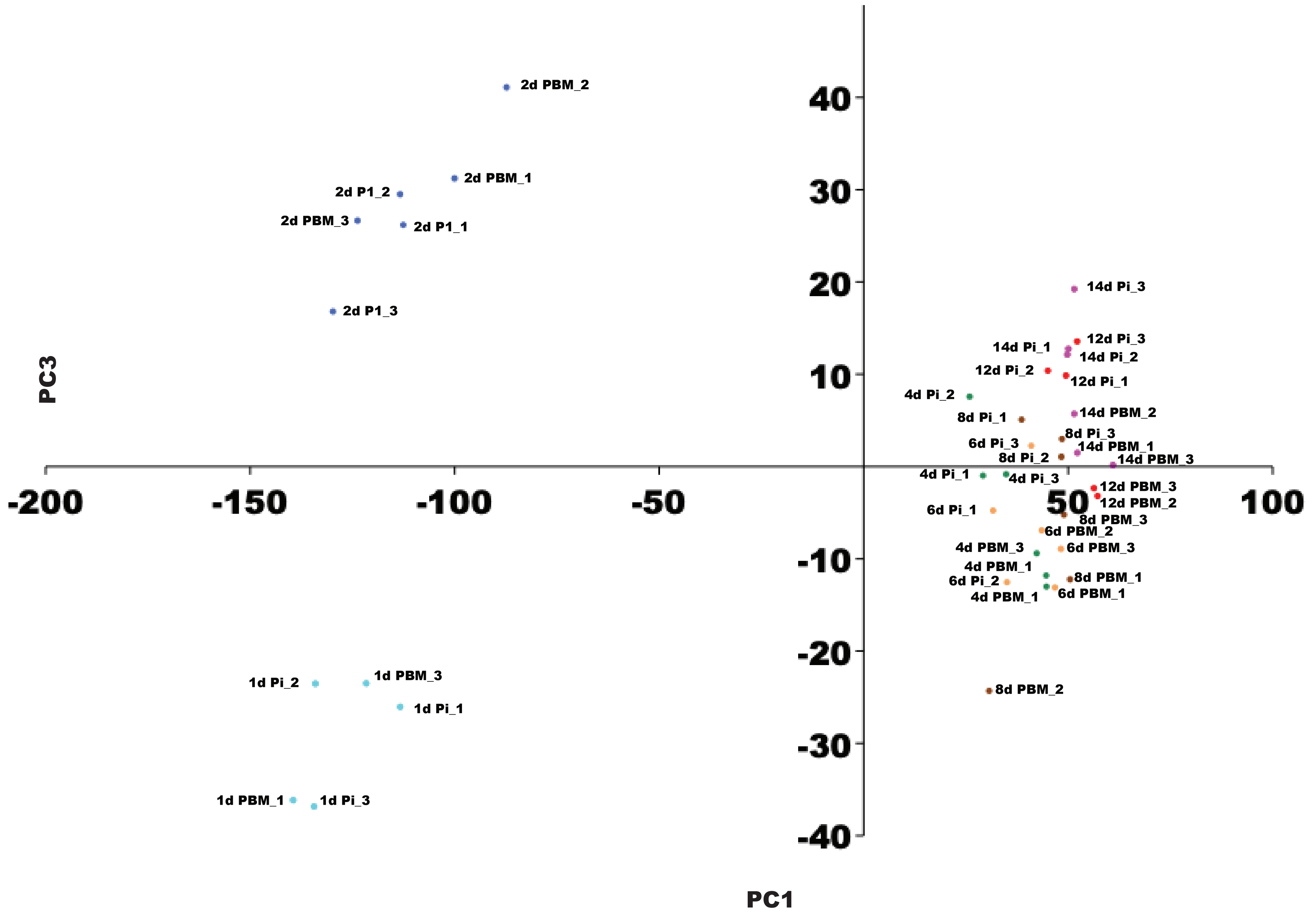

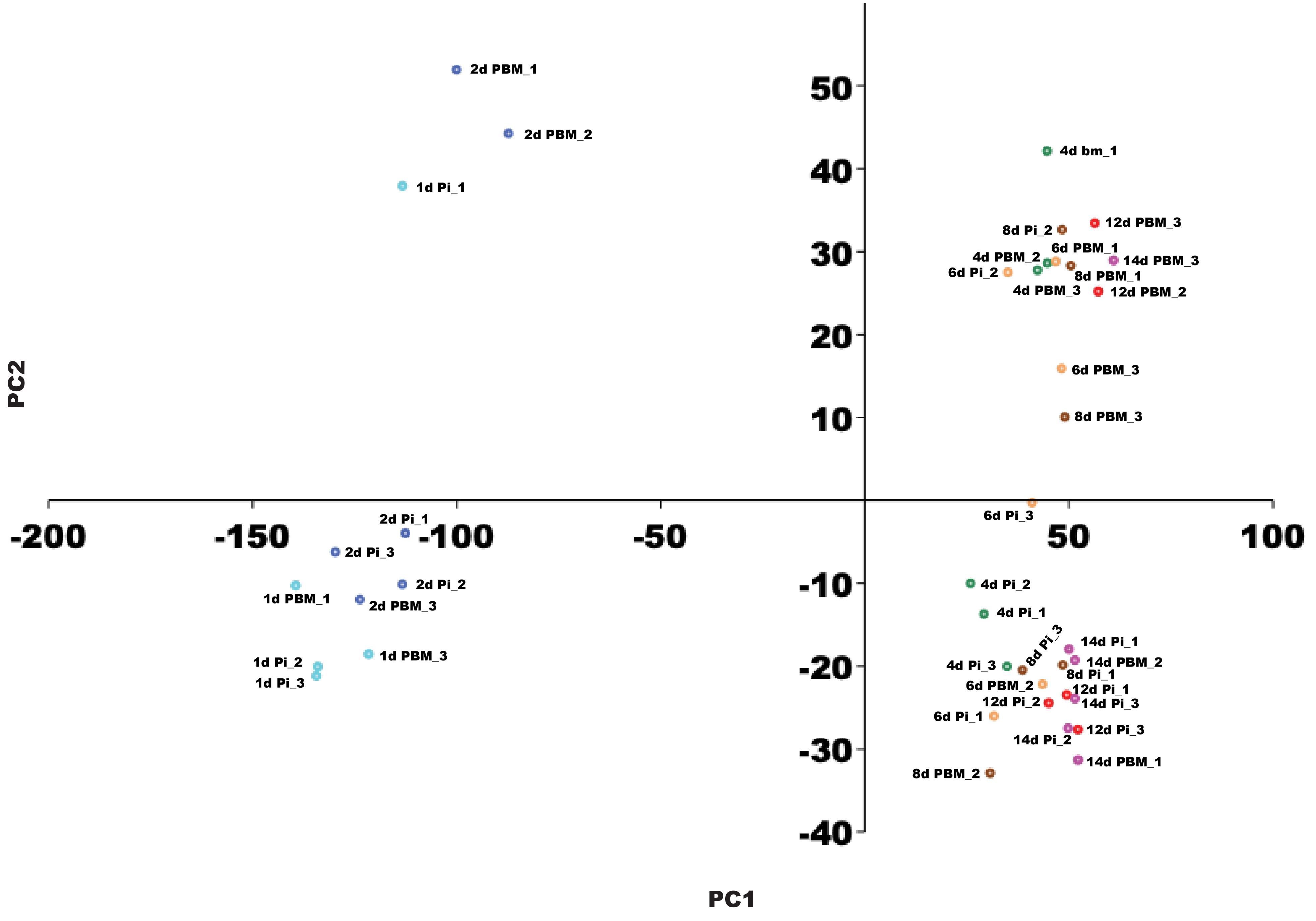

### Fig S4

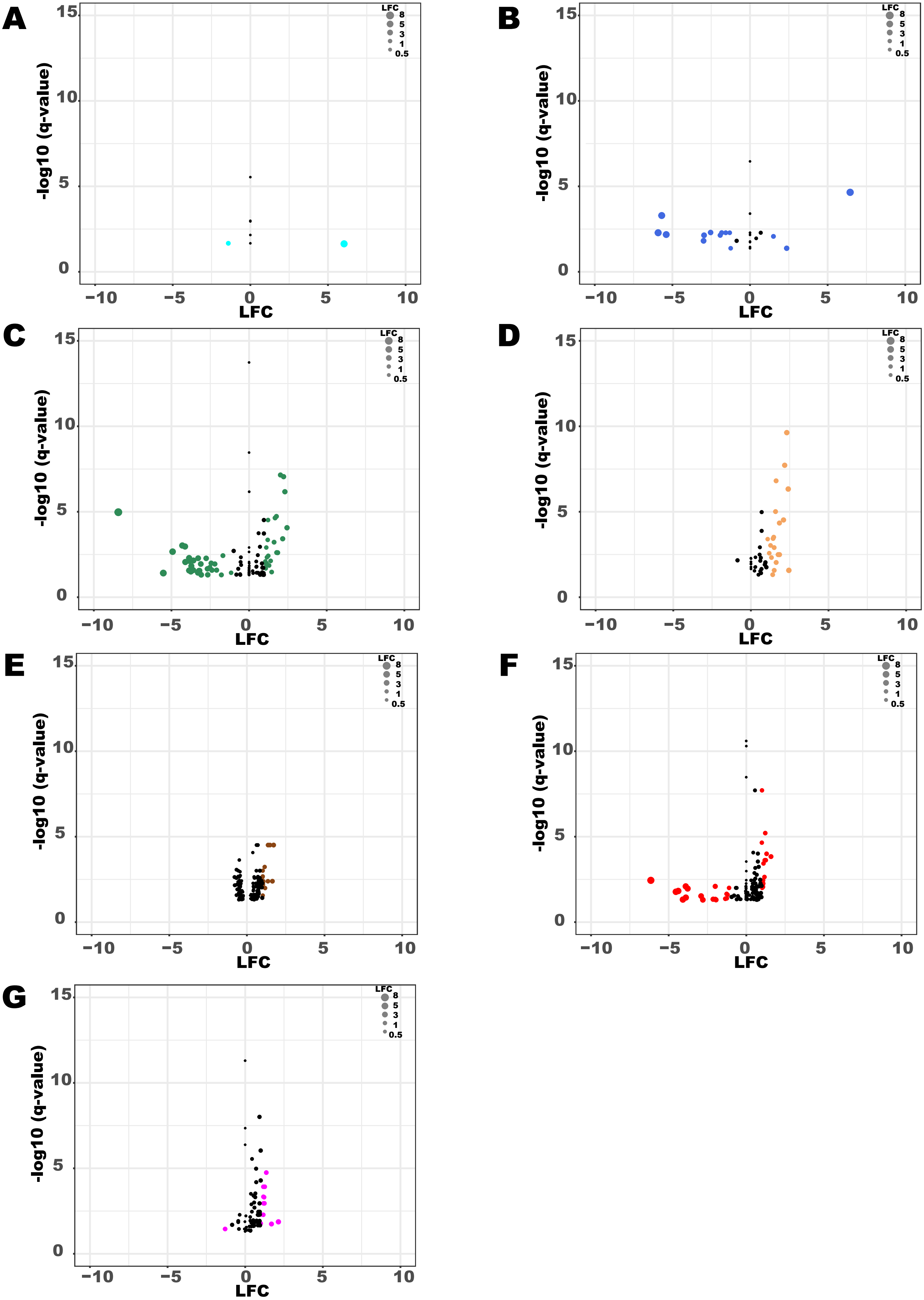

### Fig S5

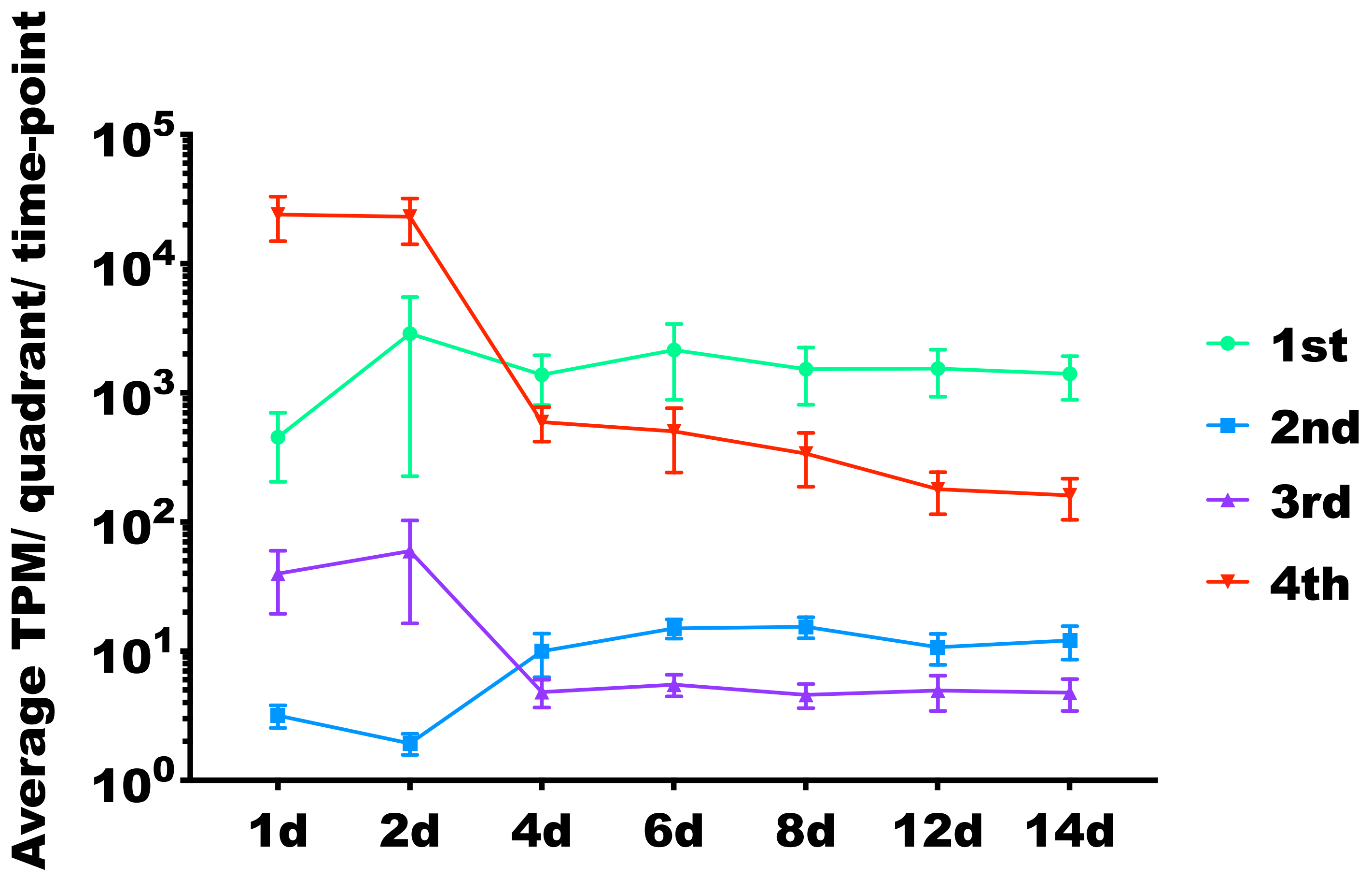
